# Individual differences in fear acquisition: Multivariate analyses of different Emotional Negativity scales, physiological responding, subjective measures, and neural activation

**DOI:** 10.1101/233528

**Authors:** Rachel Sjouwerman, Robert Scharfenort, Tina B. Lonsdorf

**Author notes:** Correspondence should be addressed to: Dr. Tina B. Lonsdorf Department of Systems Neuroscience University Medical Center Hamburg-Eppendorf 20246 Hamburg Germany.

## Abstract

Negative emotionality is a well-established and stable risk factor for affective disorders. Individual differences in negative emotionality have been linked to associative learning processes which can be captured experimentally in fear conditioning paradigms. Literature suffers from underpowered samples, suboptimal methods, and an isolated focus on single questionnaires and single outcome measures. Here, we apply multivariate and dimensional approaches for three commonly investigated questionnaires in the field (STAI-T, NEO-FFI Neuroticism, Intolerance of Uncertainty Scale) across multiple analysis units (ratings, skin conductance, startle, BOLD-fMRI) during fear acquisition-training in two large samples (N_Study1_=356; N_Study2_=113). We investigate whether the specific or shared variance of these questionnaires is linked with CS-discrimination in specific outcome measures (Study 1). We identify a significant negative association between STAI-T and CS-discrimination in SCRs and between Intolerance of Uncertainty and CS-discrimination in startle responding. Yet, correlation coefficients for all questionnaire-outcome measure combinations did not differ significantly from each other. In Study 2 the STAI-T score was positively associated with CS-discrimination in a number of brain areas linked to conditioned fear (amygdala, putamen, thalamus), but not to SCRs or ratings. Importantly, we replicate potential sampling biases between fMRI and behavioral studies regarding anxiety levels. We discuss the implications of these results.

## Introduction

Negative emotionality refers to the general tendency to show various forms of negative affect including (exaggerated) anxiety, guilt, moodiness, angriness, insecurity and dissatisfaction (Shackman, Tromp, et al., 2016). Furthermore, individuals characterized by high negative emotionality report enhanced distress in response to novelty, threat, or stress in real-life and in the laboratory (Guo et al., 2015; Michael P. Hengartner et al., 2017; Shackman et al., 2018; Shackman, Tromp, et al., 2016) as well as in the absence of real danger (Grupe & Nitschke, 2013). Negative emotionality is used synonymously with other terms in the literature as Neuroticism, Negative Affectivity and Dispositional Negativity. Here, we use the term ‘negative emotionality’ to refer to the broad umbrella construct.

Negative emotionality is a well-established and relatively stable risk factor for the development of affective disorders – in particular anxiety and depression (Barlow et al., 2014; Goldstein et al., 2018; Hakulinen et al., 2015; M. P. Hengartner et al., 2016; Michael P. Hengartner et al., 2018; Jeronimus et al., 2016; for a review see Shackman, Tromp, et al., 2016; Wichstrøm et al., 2018). Even within patient samples, those individuals with higher levels of negative emotionality typically report more severe symptoms, and clinical prognosis is less optimistic (Bufferd et al., 2018; Spinhoven et al., 2016; Steunenberg et al., 2010). To allow for the development of targeted prevention and intervention approaches, there is an urgent need to deepen our understanding on the basic neuro-cognitive mechanisms (Shackman, Tromp, et al., 2016; Shackman & Fox, 2018) underlying the elevated risk associated with negative emotionality.

Differential vulnerability might hinge on individual differences in associative learning processes (Duits et al., 2015; S Lissek et al., 2005; Mineka & Oehlberg, 2008). Associative learning processes represent a core mechanism of the development as well as the maintenance of pathological fear and anxiety. These processes can be captured experimentally in fear conditioning paradigms, which serve as translational models in fear and anxiety research (Forcadell et al., 2017; Jan Haaker et al., 2019; Scheveneels et al., 2016). During fear acquisition training, an initially neutral stimulus (the to-be-conditioned stimulus, CS+) is paired with an aversive event (the unconditioned stimulus, US) and thereby becomes a predictor of the US while a second stimulus (CS-) is never paired with the US. Subsequently, the CS+ elicits (anticipatory) defensive responses that can be assessed at different response levels, all capturing slightly different time-windows and sub-processes (for a review see Lonsdorf et al., 2017). These include self-report (e.g., ratings of fear or US expectancy), physiological responding (e.g., skin conductance responses (SCRs), fear-potentiated startle responses (FPS) and neuro-functional activation (e.g., BOLD fMRI). Skin conductance responses are the most commonly used measures of conditioned responding and are assessed as phasic arousal-related changes in sweat gland activity (Boucsein et al., 2012; Dawson et al., 2007). Fear potentiated startle, which follows a valence gradient in responding (Kuhn et al., 2019; Lang et al., 1990), measures the increase in the startle reflex elicited by a sudden event (such as a burst of white noise) in the presence of threat as compared to the absence of threat.

Focusing on individual differences in negative emotionality (e.g., Shackman, Stockbridge, et al., 2016; Shackman, Tromp, et al., 2016) and combining it with fear conditioning research (Lonsdorf & Merz, 2017) holds promise to provide critical insights into the mechanisms underlying individual risk and resilience for the development of anxiety and/or stress-related disorders (Lonsdorf & Merz, 2017; Mineka & Oehlberg, 2008). A recent review identified three scales linked to the broader construct of negative emotionality that have been consistently associated with individual differences in fear conditioning performance (Lonsdorf & Merz, 2017) and vulnerability to pathological fear and anxiety: the trait anxiety scale of the Spielberger’s State-Trait Anxiety Inventory (STAI-T (Spielberger et al., 1983)), the Big Five neuroticism scale of the NEO-FFI (NEO-FFI-N (McCrae & Costa Jr., 2004)) and the intolerance of uncertainty scale (IUS; (Gerlach et al., 2008b)).

**Trait-anxiety**, reflects the general tendency to react anxiously and to show cognitive as well as affective styles related to pathological anxiety to a wide range of events and contexts. There has been a long-standing debate on whether the STAI-T is a “good” measure of anxiety. Based on confirmatory factor analytical approaches in large samples of healthy individuals (Bados et al., 2010; Balsamo et al., 2013) and patients (Balsamo et al., 2013), it was suggested that the STAI-T measures “general negative affect” rather than “measuring anxiety or depression in a strict sense”. The latter two hypothetical sub-factors had been proposed previously (Bieling et al., 1998) but lacked sufficient discriminant validity in newer work using larger samples (Bados et al., 2010; Balsamo et al., 2013).

In turn, **neuroticism**, one of the “Big-Five” constructs derived factor-analytically, reflects the tendency to show negative affect such as anger, envy, guilt, and depressed mood and to be emotionally highly reactive and vulnerable to stress (Eysenck, 1950). Neuroticism has also been described as “sensitivity of defensive distress systems that become active in the face of threat, punishment or uncertainty” (DeYoung, 2015) and is considered an established risk factor for psychopathology (Lahey, 2009). Recently, it was reported that neuroticism may be associated with experiencing more intense negative emotions, but not with the variability in experiencing negative emotions (Kalokerinos et al., 2020).

Finally, **intolerance of uncertainty** is defined as the dispositional cognitive bias to perceive and interpret ambiguous situations as threatening (Buhr & Dugas, 2002; Gerlach et al., 2008b), which has been suggested to be a possible trans-diagnostic factor contributing to maintaining affective disorders including anxiety disorders and depression (McEvoy & Mahoney, 2012; Saulnier et al., 2019). Relatedly, patients suffering from affective disorders are characterized by heightened scores on the IUS (Boswell et al., 2013). Of note, several different scales assessing intolerance of uncertainty co-exist (Fergus, 2013; Gerlach et al., 2008b).

All three constructs (trait anxiety, neuroticism and intolerance of uncertainty), as assessed through the above mentioned scales, are related to – or can be subsumed under – the broader umbrella negative emotionality. All three have been associated with individual differences in fear conditioning performance (Lonsdorf & Merz, 2017). Yet results in the literature are heterogeneous and partly inconclusive at the behavioral and neuro-functional level. The fear conditioning field, similar to the field of personality neuroscience in general (Allan & DeYoung, 2016), suffers from a number of well described problems. These problems include: **(1)** generally underpowered samples (typically below N=30 per group (Lonsdorf & Merz, 2017), **(2)** sub-optimal statistical approaches such as dichotomizing continuous variables which gives rise to interpretation problems, causing massive loss of power and increases in Type II error rates (i.e., false negatives) (Altman & Royston, 2006; Cohen, 1983; McClelland & Irwin, 2003; Preacher et al., 2005). Furthermore the majority of results in the fear conditioning field originate from univariate analyses focusing on **(3)** single constructs related to negative emotionality (for a discussion see Lonsdorf & Merz, 2017), for a few exceptions see (Chin, Nelson, Chin, et al., 2016; Dunsmoor et al., 2016; Morriss et al., 2015, 2016; Otto et al., 2007) and **(4)** singular outcome measures (such as ratings, SCRs, FPS or BOLD fMRI) each tapping into slightly different underlying processes (Lonsdorf et al., 2017). Attempts for multivariate integration are thus far rare. As a consequence, separate lines of research and isolated findings have emerged that are notoriously difficult to integrate and interpret into one bigger picture. Hence, we echo recent calls for a paradigm shift embracing more complex multivariate approaches, the use of larger data sets and the dimensionality of the data (Holmes & Patrick, 2018; Shackman & Fox, 2018). The overarching aim of this work is to enhance our understanding of the mechanisms through which negative emotionality may convey risk for affective psychopathology by integrating separate lines of research (using different scales and outcome measures) that have emerged in parallel and are difficult to integrate.

To achieve this aim, we start by integrating dimensional measures as derived from three commonly employed scales in the field (i.e., STAI-T, NEO-FFI-N and IUS) with the three most commonly used measures of conditioned responding (ratings, SCRs, FPS) – as identified and summarized by a recent review by our group (Lonsdorf & Merz, 2017). These measures are obtained in a large sample (**Study 1**, N=356) and combined into one statistical model that is set up to investigate whether any of these scales is linked to specific fear conditioning performance over-and-beyond the other scales. Subsequently we investigate whether it is the shared variance across the scales and across outcome measures that explains these potential associations and thus supports a prominent role for the general construct negative emotionality, or whether the scales remain specifically associated with specific measures of conditioned responding. Specific and directed hypotheses on the outcome of this model are difficult to derive from the existing literature because results in the field are extremely heterogeneous.

Additionally, we aim to replicate the main findings from Study 1 in a re-analysis of a second pre-existing sample (**Study 2**, N=113) which also allows to extend our inferences to the neuro-functional level (a brief Introduction to Study 2 is provided below).

### Study 1: Methods

#### Participants

Three-hundred-fifty-six healthy individuals participated in Study 1. The study was originally designed to investigate individual differences in fear acquisition and to investigate post-acquisition manipulations on return of fear responding (see also experimental design). Participants were recruited primarily through online advertisement on a local student-job website.

Prior to inclusion, individuals were subject to a pre-experimental telephone screening in which individuals were selected only when reporting absent previous or current diagnosis of psychiatric or neurological disorders, or hormonal disturbances such as thyroid dysfunction. This sample includes 255 females and 100 males between 18 and 40 years old, with an average age of 25 ± 4 (SD) years. Note that gender information is missing for one participant. Written informed consent in accordance with the Declaration of Helsinki was obtained from each participant, and the Ethical Review Board of the German Psychological Association (DGPS) approved the study. Participants received 10 Euros/h for their participation. Please note that this sample and the association between differential SCRs during fear acquisition training and scores on the STAI-T scale (see below) has been included as a case example in our recent publication focusing on the impact of performance-based exclusion of participants (i.e., exclusion based on differential SCRs cut-offs) to illustrate a potential sample bias with respect to individual differences in anxiety related traits (Lonsdorf, Klingelhöfer-Jens, Andreatta, Beckers, Chalkia, Gerlicher, Haaker, et al., 2019).

#### Questionnaires

Participants filled in a batch of questionnaires prior to the experiment. This batch included 1) questions to obtain demographic information, 2) the State-Trait Anxiety Inventory (Spielberger et al., 1983), 3) the NEO-FFI (Gerhard, 1999; McCrae & Costa Jr., 2004) 4) the Intolerance of Uncertainty Scale (Gerlach et al., 2008a) and 5) the IPC (Krampen, 1981). Upon completion of the experiment (i.e., after extinction and reinstatement), participants filled in a post-experimental awareness questionnaire (Lonsdorf et al., 2017) of which answers were orally confirmed with the experimenter. The questionnaire included estimations on the total number of received electrotactile stimuli and the total number of experimental stimuli presented during the experiment. Also, it contained questions about perceived CS-US contingencies during the experiment (first as a free recall then as a forced choice). Based on this, participants were classified as either aware (N=236, able to correctly report CS-US contingencies in free recall and/or forced choice) or unaware (N=87, unable to report correct CS-US contingencies across questions). Twenty-one participants that reported a tendency towards the correct contingencies but also some unsureness were counted as aware. Data on CS-US awareness were missing from twelve participants.

The trait scale of the STAI (STAI-T) consists of 20 items, evaluated on a four-point Likert scale, allowing individuals to score between minimally 20 and maximally 80 points. Despite its potential misleading name (i.e. trait anxiety inventory), the STAI-T more likely assesses how a respondent generally feels, and is thought to target relatively stable aspects significant for “anxiety proneness”, including calmness, confidence and security (Julian, 2011). Congruently, the STAI-T has been criticized for representing a psychometrically inhomogeneous scale itself (Bados et al., 2010; Balsamo et al., 2013) representing facets of anxiety and depression. Based on confirmatory factor analytical approaches in large samples of healthy individuals (Bados et al., 2010; Balsamo et al., 2013) and patients (Balsamo et al., 2013), it was suggested that the STAI-T measures “general negative affect” rather than “measuring anxiety or depression in a strict sense”. The latter two hypothetical sub-factors had been proposed previously (Bieling et al., 1998) but lacked sufficient discriminant validity in newer work using larger samples (Bados et al., 2010; Balsamo et al., 2013).

The neuroticism scale of the NEO-FFI (NEO-FFI-N) consists of 12 out of 60 items, which were derived factor analytically and should represent one of the five higher order Big-Five personality traits (McCrae & Costa Jr., 2004), i.e., neuroticism. Scores on this NEO-FFI-N scale can range between 0 and 48. Neuroticism refers to the tendency to express negative emotionality and has been suggested to be associated with defensive responding to uncertainty, threat, and punishment (Allen & DeYoung, 2017).

The IUS consists of 27 items that aim to assess an individual’s tendency to react to the uncertainties of life, or more precisely their intolerance towards these uncertainties (Buhr & Dugas, 2002; Gerlach, Andor, & Patzelt, 2008). Each item is evaluated on a five-point Likert scale, allowing respondents to score between 27 and 135. Intolerance of uncertainty is defined as a cognitive bias that affects how uncertain situations are perceived, interpreted, and dealt with (cf. Dugas et al., 1998; Freeston et al., 1994). Several factor solutions have been suggested including a four-factor solution (Buhr & Dugas, 2002) which include: 1) uncertainty is stressful and upsetting, 2) uncertainty causes inability to act, 3) uncertain events are negative and should be avoided, and 4) being uncertain is unfair. No official German version of this questionnaire exists, however, Gerlach and colleagues (Gerlach et al., 2008c) created a German translation and investigated the underlying factor structure in which this four-factor structure could not be replicated. In Study 1, this German translation of the full 27 items of the IUS is used.

For those individuals having one or more, but not all, missing items on either of the questionnaires, missing values were imputed using a single imputation with the predictive mean matching method in the MICE package in R. This imputation method draws observed values from other subjects with a similar response pattern on other variables. In total, data was imputed for 8 subjects on the STAI-T, for 5 subjects on the NEO-FFI-N, and for 26 subjects on the IUS. Forty-six participants have missing data for the full STAI-T and IUS, one misses the full STAI-T only, and one participant has full missing data on the NEO-FFI-N. Because this data cannot be considered as missing at random, it is not imputed, but maintained as missing data.

Overall reliabilities of the questionnaires and subscales of interest were high in the final sample, as indicated by Cronbach’s α: 0.91 for STAI-T; 0.86 for NEO-FFI-N; 0.94 for IUS. In addition, a wide range of scores was covered in the acquired sample: 21-76 for STAI-T with 38 ± 9 (mean ± sd); 1-40 for NEO-FFI-N with 20 ± 8 (mean ± sd); and 27-135 for IUS with 62 ± 18 (mean ± sd).

#### Experimental design

All participants underwent a fear conditioning, extinction and return of fear paradigm. Data of interest for the current research question concerns fear acquisition training only. Data acquired during the same experimental session but involving experimental manipulations after this fear acquisition training phase or involving a methodological validation in sub-samples of this sample is published elsewhere (Sjouwerman et al., 2015; Sjouwerman & Lonsdorf, 2018). Therefore, experimental details will only be provided for the fear acquisition training phase and the preceding habituation phases of the experiment.

#### Instructions

Participants were not instructed with respect to the CS-US contingencies or the learning element of the study.

#### Visual material – conditioned stimuli

Black geometrical shapes (i.e., a rectangle and an ellipse) served as conditioned stimuli (CS). One of these shapes (CS+) co-terminated with the unconditioned stimulus (US) during all fear acquisition training trials, whereas the other shape did not (CS-). In other words, a 100% reinforcement ratio was used during this experimental phase. Each CS type was presented consecutively for maximally two times, and nine times in total during fear acquisition training (9 CS+ and 9 CS-trials). Allocation of the shapes to CS+ or CS-was counterbalanced across participants, as well as the order in which the CS+/CS− appeared. The CSs were presented for 6 seconds on a colored computer screen (blue, purple, green, or yellow). The background color served as contextual stimulus, which has no relevance to the fear acquisition training phase, but is of value in the context of post-acquisition experimental manipulations – not included here. The background color remained constant within experimental phases and was counterbalanced across participants. CS presentations were interleaved with inter trial intervals (ITI) consisting of a white fixation cross on a black computer screen, with variable durations (11.5 ± 1.5 seconds). Prior to fear acquisition training, subjects underwent an explicitly US-free CS habituation phase in which both stimulus types (i.e., the CS+ and CS-) were presented two times each.

#### Electro-tactile material – unconditioned stimulus

A train of three electro-tactile square wave pulses, 2 milliseconds each, with 50 millisecond intervals, served as US. The US was produced by a DS7A electrical stimulator (Digitimer, Welwyn Garden City, UK) and delivered through a surface electrode with a platinum pin (Specialty Developments, Bexley, UK) to the dorsal part of the right hand. The intensity of the electro-tactile US was individually adjusted using a stair-case procedure to reach an unpleasant but tolerable level (range US intensities: 0.3-70mA, mean ± SD = 4.7 ± 5.2, median = 3.5). The intensity of the US was gradually increased after conferring with the participant. The participant could then herself/himself elicit the US by pressing the space bar. After delivery of each US, the participant rated the averseness of the US on a scale from one to ten, with one not being aversive at all to ten being unbearable. The experimenter aimed to reach a final averseness rating of seven, which was not explicitly communicated to the participant.

#### Acoustic material – startle probes

A burst of 95dB(A) white noise was used to elicit a startle response. Startle probes were presented binaurally via headphones (Sennheiser, Wedemark, Germany) four or five seconds after CS onset in half of all CS habituation trials (one out of two trials) and in two thirds of all CS fear acquisition training trials (six out of nine trials). Additionally, startle probes were presented in one third of all ITI’s, either five or seven seconds after ITI onset. To obtain a stable baseline for startle reactivity, five consecutive startle probes – interleaved six seconds – were presented during a white fixation cross on a black computer screen.

#### Procedure

Experimental instructions were provided in written form and importantly did not contain instruction with respect to CS-US contingencies. Participants were instructed to attend to the visual stimuli presented on the screen, and ignore the acoustic startle probes. It was made explicit that startle probes had the sole purpose of enabling physiological data acquisition.

Participants started by filling out the questionnaires. After, they proceeded with the US intensity calibration phase. In a step-wise procedure the US intensity was increased to a level described by the participant as very annoying but not painful equaling to a rating of at least 7 on a ten-point scale (with ten being the maximally aversive sensation that could be induced by the electrode). Next, the actual experiment started with the startle habituation phase and continued with an explicitly US-free CS habituation phase. Subsequently the uninstructed fear acquisition training phase of interest started. Presentation of all stimuli was controlled using Presentation Software (NeuroBehavioral Systems, Albany California, USA). After completing the full experiment, thus after the post-acquisition training phases that included extinction training, reinstatement administration, and return of fear test phases, participants completed the post experimental awareness questionnaire. Twenty-eight participants had to be excluded for the fear conditioning experiment either due to voluntarily discontinuation or technical failure during data acquisition.

#### Subjective data recording – Fear ratings

Participants indicated their level of fear, anxiety, and distress towards both CS types within intermittent rating blocks. The following text was presented on screen: “How much stress, fear or anxiety did you experience the last time you saw symbol X?”, with the “X” referring to one of the CS types at a time. Participants were given seven seconds to provide their response on the computerized visual analogue scale (VAS) ranging from 0 (none) to 100 (maximum), which had to be confirmed within the given time window by pressing the enter key. One rating block was presented at the end of the habituation phase, and three rating blocks were presented during fear acquisition training. Rating blocks were always presented after minimally one and maximally four CS+ and CS-presentation(s) (see (Sjouwerman et al., 2016a) for a graphical overview on the design including ratings). The last rating in the fear acquisition training phase either occurred after the seventh or eighth acquisition trial. Nineteen participants failed to confirm their selected VAS values within seven seconds, and have therefore missing rating data. Post processing was conducted in R version 3.6.0 (2019-04-26).

#### Physiological data recording and processing – Skin conductance and startle responding

Physiological data were recorded using a BIOPAC MP100 amplifier, (BIOPAC Systems Inc., Goleta, CA) and AcqKnowledge 3.9.2 software. Data preprocessing was conducted in MATLAB (version2014b), response quantification was conducted manually in a custom made program, and post processing was conducted in R version 3.6.0 (2019-04-26). For physiological measurements, additional data is missing for some participants (SCR = 8, startle = 30) due to technical failures including for example saving failure of the physiological data file only, data extraction problems, erroneously adjusting the gain during the experiment causing SCR amplitudes to be uninterpretable, or electrode misplacement during data acquisition.

For **skin conductance** recording, participants first cleaned their hands with warm water. After, two hydrogel and Ag/AgCl sensor recording electrodes (Ø 55 mm) were attached to the distal and proximal hypothenar eminence of the left hand. Skin conductance data were recorded continuously at 1000 Hz with a gain of 5mΩ. In case participants’ skin conductance moved beyond the scaling window, however, the gain (i.e. resistance) was increased or decreased to reduce or increase sensitivity of the skin conductance being recorded prior to the start of the experiment. Offline, data was down sampled to 10Hz. According to published guidelines (Boucsein et al., 2012), data were scored manually as foot-to-peak responses with response onsets starting between 0.9 and 4.0 seconds after CS or US onset. Increases smaller than 0.02 µS were scored as zero responses. Responses confounded by recording artifacts, such as electrode detachment, responses moving beyond the sampling window, or excessive baseline activity were discarded and scored as missing values. Raw skin conductance response (SCR) amplitudes were normalized by log transformation and range corrected by division through an individuals’ maximum response amplitude (either CS or US).

#### Physiological SCR non-responder

Participants not showing valid SCRs in over two thirds of all fear acquisition training to the US (i.e., six out of nine trials (Lonsdorf, Klingelhöfer-Jens, Andreatta, Beckers, Chalkia, Gerlicher, Haaker, et al., 2019)) were classified as physiological non responder (n = 16) and all SCR trials were set to missing values.

**Startle responding** was measured by using Ag/AgCl electromyogram (EMG) electrodes. Two electrodes were placed below the right eye over the orbicularis oculi muscle and one electrode was placed on the participants’ forehead to obtain a reference signal. Startle data filtered online (band-pass: 28-500 Hz), rectified, and integrated (averaged over 20 samples). According to published guidelines (Blumenthal et al., 2005) data were scored manually as foot-to-peak with response onsets within 20-120 milliseconds post startle probe onset. Responses confounded by a blink occurring up to 50 milliseconds before startle probe onset were scored as missing value. Similarly, trials confounded by recording artifacts or excessive baseline activity within the same time window were scored as missing values. Raw data were t-transformed across the experimental phases up to the fear acquisition training phase.

#### Physiological startle non-responder

Participants not showing valid startle responses in over one third of all trials from the habituation phases and the fear acquisition training (i.e., more than 9 out of 28) were classified as physiological non responder (n = 16), and startle responses for these participants were set to missing values.

Fear acquisition is operationalized as CS+/CS− discrimination during the fear acquisition training phase (i.e., average CS+ minus average CS-responding). This includes responding towards 9 CS+ and 9 CS-trials for SCRs, and 6 CS+ and 6 CS-trials for startle responses (not all trials were ‘startled’, see above), and all 3 intermittent CS+ and 3 intermittent CS-ratings.

#### Study 1: Statistical analyses

Study 1 consists of three analysis steps. **First**, univariate zero-order correlational analyses are conducted between the three questionnaires, or their respective subscale, and the three outcome measures of conditioned responding recorded in our study. All variables are treated as dimensional. For all univariate analyses, multiple testing will be corrected by using the Benjamini Hochberg method (p_BH_). Correlation coefficients were compared according to (Lee, I.A., 2013).

In a **second step** which serves the aim to integrate potential effects of these independent and dependent measures in a single model, a path model is constructed in which relationships between the three independent variables and the three dependent variables are estimated simultaneously. In this path model, correlations among the independent measures, i.e., questionnaires or scales, are allowed.

In a **third step** we take into account that the three independent measures are likely to highly correlate with each other which makes it possible that these questionnaires, or subscales of questionnaires, all tap into a same larger construct, i.e., “negative emotionality”. Similarly, the three measures of conditioned responding highly correlate and might be part of the larger construct “fear learning”. This hypothesis, i.e., whether it is the shared variance across the questionnaires or scales or unique variance of individual questionnaires or scales that explains differences in specific outcome measures will be examined by employing a structural equation model. In this model, two latent variables will be defined, one for the three questionnaires/scales and one for the three outcome measures. The regression weight between the latent variable negative emotionality and the first questionnaire/scale, as well as the regression weight between the latent variable fear learning and the first outcome measure will be fixed to 1. Subsequently, a complementary structural equation model will be constructed that in addition to the paths defined in the model described above (under step 3) includes the paths that showed significant associations in the established path model in step 2.

For both, path models and structural equation models, two-sided model fit will be evaluated based on root mean square error of approximation (RMSEA) values, indicating excellent fit when < 0.01, good fit when < 0.05, fair fit when < 0.08, mediocre fit when < 0.10, and poor fit at > 0.10 RMSEA values (M. W. Browne, 1992; MacCallum et al., 1996). To improve model fit, backward selection of significant and trend-significant path will be executed. Trends (p < 0.1) will be included in interim models, but not in final models. Full models, interim models, as well as final models will be reported.

Univariate statistical analyses and data visualization was performed with R version 3.6.0 (2019-04-26) using packages corrplot, dplyr, ggplot2, tydr, mice, psych and cowplot. Multivariate analyses (path and structural equation models) were performed in AMOS 26 for Windows (Armonk, NY).

### Study 1: Results

Study 1 aims to enhance our understanding of the mechanisms through which negative emotionality may convey risk for affective psychopathology by integrating separate lines of research (using different scales and outcome measures) that have emerged in parallel and are difficult to integrate. In a **first step**, we present univariate analyses illustrating associations between the three commonly employed scales in the field (i.e., STAI-T, NEO-FFI-N and IUS) with the three most commonly used measures of conditioned responding (ratings, SCRs, startle responding). We then move to multivariate analyses integrating these variables into a single model (**Step 2**) and exploring the role of potentially latent higher-order factors (**Step 3**).

#### Step 1: Univariate analyses

Univariate analyses revealed a significant albeit small negative correlation between STAI-T and CS discrimination in SCRs (r = −0.19, p_BH_ = 0.007), whereas correlations between CS discrimination in SCRs and either NEO-FFI-N or IUS were not significant when correcting for multiple testing but were trend wise significant and significant only when not correcting for multiple comparisons (see Figure 1). This effect seems descriptively driven by the combination of weakly and non-significantly increasing CS-responding, and weakly and non-significantly decreasing CS+ responding with increasing scores on these questionnaires/scales. The correlation coefficients for CS discrimination in SCRs and the three questionnaires do however not differ significantly from each other (SCR discrimination for STAI-T and NEO-FFI-N: z = −1.691, p = 0.091; STAI-T and IUS: z = −0.977, p = 0.329; NEO-FFI-N and IUS: z = 0.498, p = 0.619).

**Figure 1.**
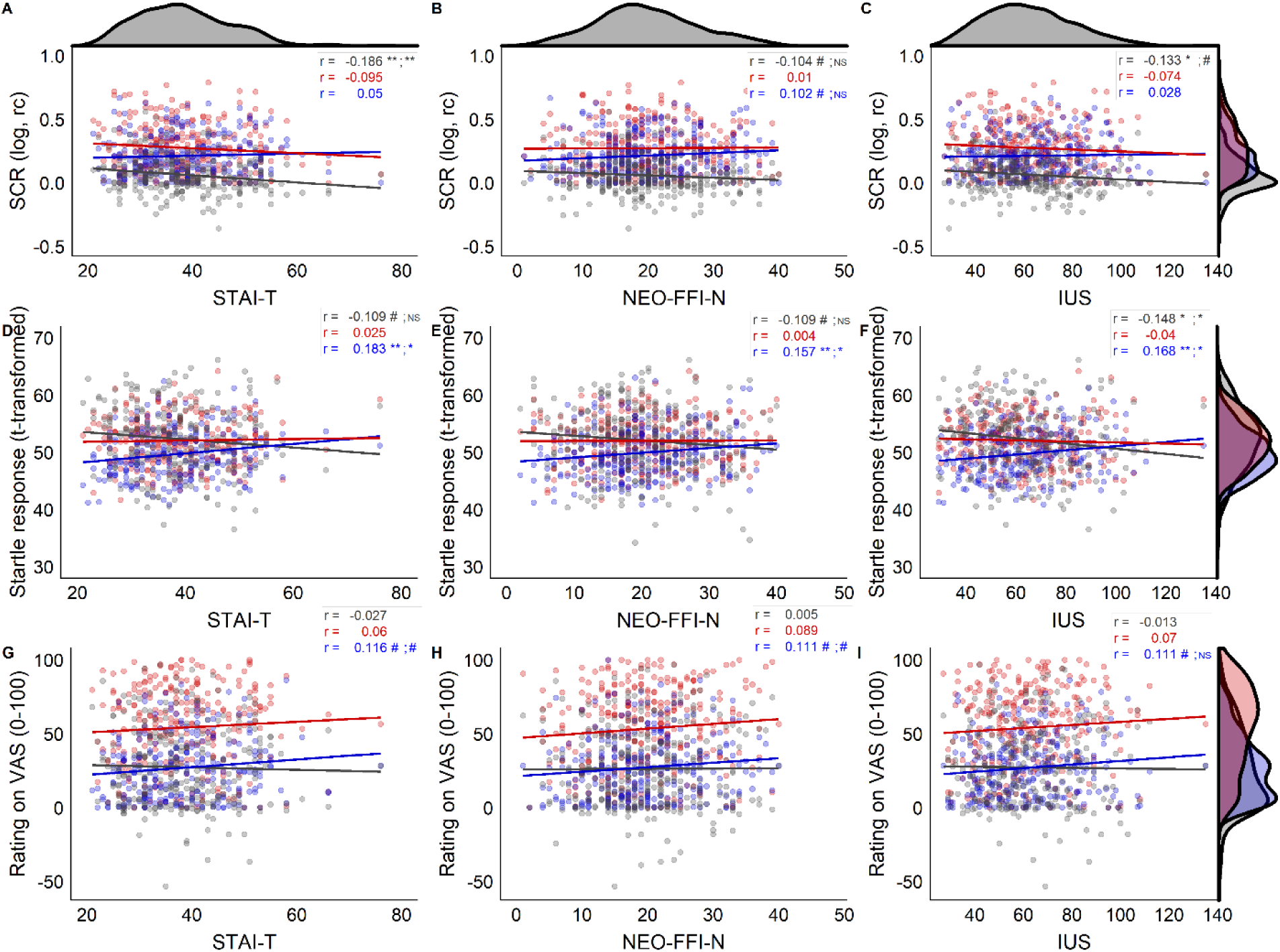
Scatterplots showing associations between the three independent variables, i.e., questionnaires/scales STAI-T (A,D,G), NEO-FFI-N (B,E,H), and IUS (C, F, I) and the three dependent variables, i.e., outcome measures SCR (skin conductance responses; A,B,C), startle responses (D, E, F), and ratings (G, H, I) for CS discrimination in grey, CS+ responding in red, and CS-responding in blue. Corresponding correlation coefficients (r) are displayed in corresponding colors, # indicates p < 0.1, * indicates p <. 05, ** indicates p < 0.01 for raw and Benjamini Hochberg (BH) adjusted p-values separated by a semicolon. NS or blank spaces indicate p > 0.1. Density distributions of scores on the questionnaires are displayed on top of the figure, density distribution per stimulus type (CS+, CS-) and for CS discrimination are displayed on the right side of the figure for each dependent variable (SCR, startle, ratings). Note that the scale of CS discrimination values for startle responding are transformed with +50 for illustrative purposes to ease comparison with CS+ and CS-startle responding.

Furthermore, a small negative correlation between IUS and CS discrimination in startle responding (r = −0.15, p_BH_ = 0.039) during fear acquisition training was observed, with no (BH corrected) significant associations for STAI-T and NEO-FFI-N and CS discrimination in startle responding but only trend wise significant associations when not controlling for multiple comparisons. The negative association between IUS and startle responding is driven by significantly increasing CS-responding with increasing scores on the IUS, while the association between the CS+ and scores on the IUS is not statistically significant. Even though the associations for the NEO-FFI-N and STAI-T with CS discrimination were uncorrected only trend significant, a similar pattern (i.e., significant positive association; r = 0.157 – 0.183, p_BH_ < 0.05) between these questionnaire scores and CS-responding was observed. The correlation coefficients for the CS discrimination and each of the three questionnaires/scales were not significantly different (all z < 0.69, all p > 0.492).

CS discrimination in ratings was not significantly associated with any of the three questionnaires, or scales of questionnaires. But note that CS-responding in ratings, similar to the pattern observed in startle responding, was positively – albeit only trend wise - associated with all three questionnaires/scales.

#### Exploratory analyses testing for an association with the STAI-*T* score and US intensity as well as awareness

As previous research has suggested an association between awareness and US intensity with trait-anxiety (for a review see Lonsdorf & Merz, 2017) exploratory analyses were conducted for Study 1 to exclude that results are biased by these factors. In brief, neither awareness of CS contingencies nor US intensity were significantly associated with the STAI-*T*: US intensity in mA did not correlate with STAI-*T* scores (r = −0.06, *p* = 0.26). Individuals aware (n = 236, mean STAI-*T* = 38) unaware (n = 87, mean STAI-*T* = 40) or uncertain (n = 21, mean STAI-*T* = 39) of CS-US contingencies did not differ significantly in STAI-*T* scores (F(3,297) = 1.51, *p* = 0.21).

#### Step 2: Multivariate analyses

Path analysis on the sum scores on the three questionnaires/scales and the three outcome measures of fear learning revealed the expected significant positive paths between all questionnaire scores and between all outcome measures (all p’s < 0.001). In addition, a significant path between STAI-T and CS discrimination in SCRs was observed in this full model in which all paths between all variables were included (Figure 2, grey font and grey paths). The final model generated through backwards selection (Figure 2, blue font and blue path), also yielded a significant path between IUS and CS discrimination in startle, which was only trend-significant in the full model.

**Figure 2.**
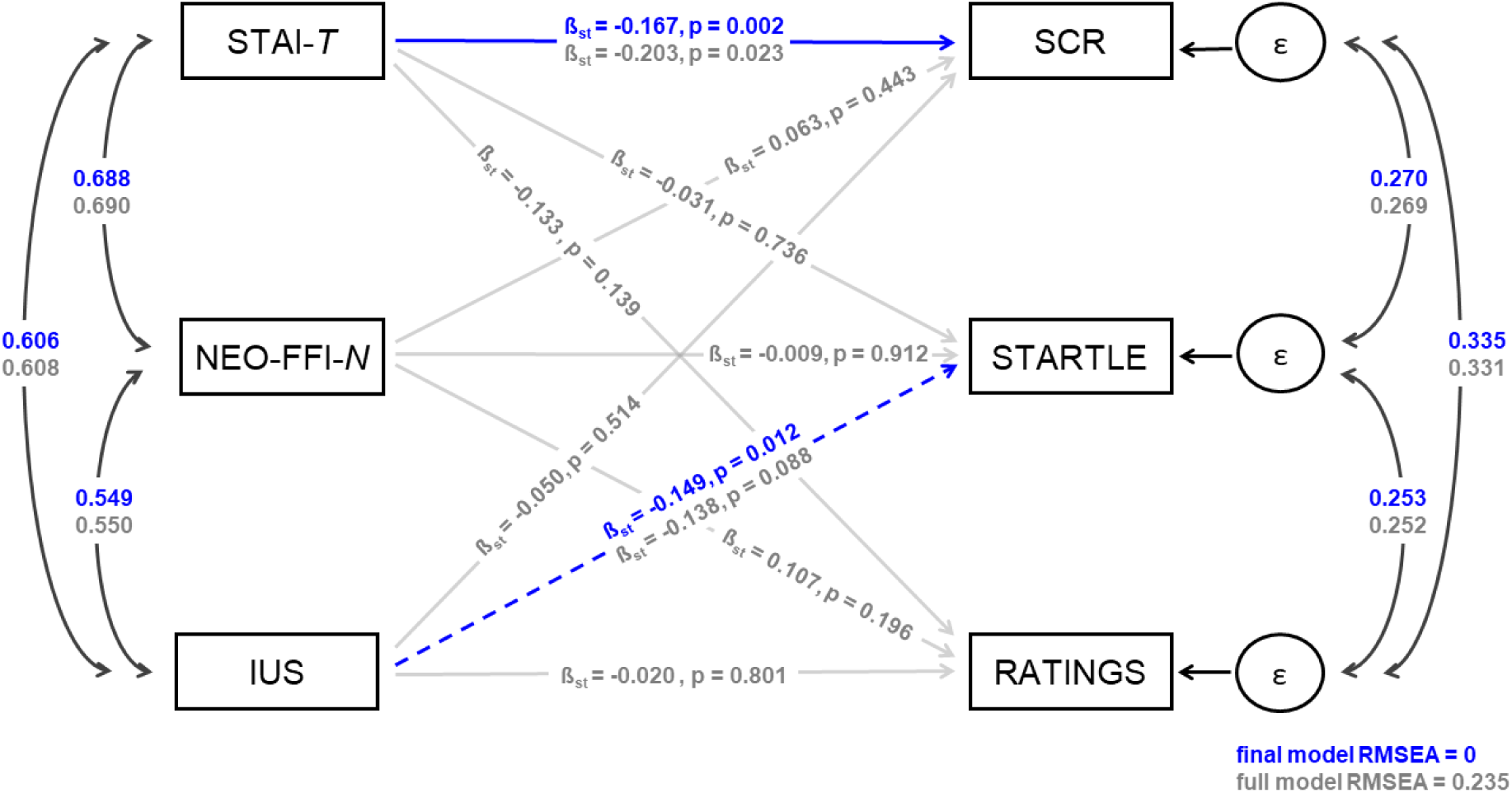
Path model reflecting the associations between the three measured questionnaires or scales of questionnaires (STAI-T, NEO-FFI-N, and IUS) with three outcome measures of fear acquisition (CS+, CS- discrimination in SCR, STARTLE, and RATINGS). Path are labeled with standardized regression coefficients (ß_st_) and p-values. Full model paths and coefficients are shown in grey; paths and corresponding coefficients selected in the final model are overlaid in blue. Note that solid grey lines represent non-significant path (p > 0.1) in the full model. Blue dashed lines represent path that were trend significant (p < 0.1) in the full model, and significant in the final model (p < 0.05), solid blue lines represent path significant (p < 0.05) in the full model and significant in the final model. Correlation path between independent and between dependent measures are shown in black, all p < 0.001 unless otherwise specified. Coefficients appear in grey for the full model and in blue for the final model. The full model shows poor fit (RMSEA > 0.10) while the final model shows excellent fit (RMSEA <0.01).

#### Step 3: Multivariate analyses testing for shared vs. unique variance

To test the hypothesis whether the associations observed in step 2 are specific to specific questionnaire scores or driven by shared variance of the three questionnaires or scales, we set up three structural equation models: (1) an initial model with two latent variables (“negative emotionality” and “fear learning”, Figure 3, grey lines and font), (2) an interim model (Figure 3, black lines and font) in which we added the unique paths identified in Step 2 to the initial model (i.e., STAI-T and SCR discrimination as well as IUS and startle discrimination), and (3) a final reduced model generated through backward selection (Figure 3, blue lines and blue font).

**Figure 3.**
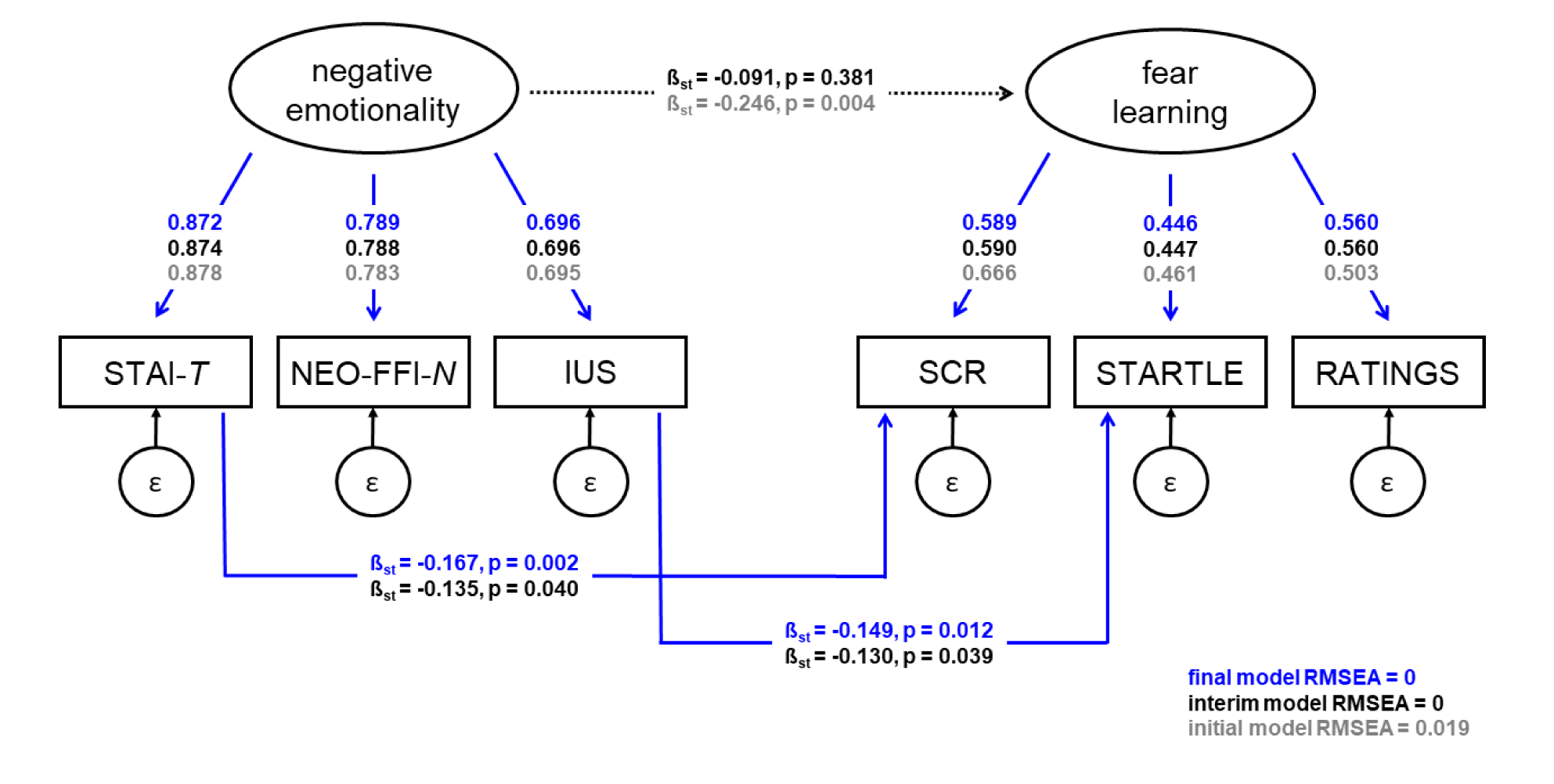
Structural equation model reflecting the relation between two latent factors: “negative emotionality” comprised of the three measured questionnaires/scales STAI-T, NEO-FFI-N, and IUS, and “fear learning” comprised of the three outcome measures of fear learning (CS+, CS-discrimination in SCR, STARTLE, and RATINGS). Path are labeled with standardized regression coefficients (ß_st_) and p-values. Initial model coefficients are shown in grey; paths and corresponding coefficients selected in the final model are overlaid in blue. Path and coefficients of the interim model are shown in black, note that the dotted black line reflects a path from an interim model significant in the initial model, but non-significant (p > 0.1) when adding specific questionnaire to outcome path, and thus is excluded for the final model. Factor loadings between latent and respective independent and dependent measures appear in grey for the initial model, in black for the interim model and in blue for the final model. All p < 0.001 unless otherwise specified. All p < 0.001 unless otherwise specified. The initial model shows good fit (RMSEA < 0.05), interim and final models show excellent fit (RMSEA <0.01).

The **initial model** shows that the three questionnaires or scales, STAI-T, NEO-FFI-N, and IUS are indeed closely related to the latent variable “negative emotionality” with all factor loadings > 0.69. Similarly, the three measures of fear acquisition, SCR, startle responding and ratings are also closely related to the latent variable “fear learning” with slightly lower factor loadings (yet, all > 0.46). This pattern is maintained in the interim and final model. Importantly, in this initial model (Figure 3, grey lines and font) in which the path between STAI-T and SCR, and IUS and startle responding were not yet included, a significant negative relation (ß_st_ = −0.246) between the two latent factors “negative emotionality” and “fear learning” was observed, suggesting that there is a general predictive effect of higher negative emotionality being linked to reduced differential fear learning. This model shows good fit (RMSEA = 0.019).

In the **interim model** the two unique significant paths identified in Step 2 (i.e., STAI-T and SCR discrimination as well as IUS and startle discrimination) were added to the initial model. This resulted in the relation between the two latent factors disappearing (black dotted line in Figure 3), whereas the two unique path turn out significant in this interim model. This path structure is entered in the **final model** and the paths between STAI-T and SCR and IUS and startle remain significant. The interim and the final model both show excellent fit (RMSEA = 0). This suggests that it may be the unique variance in STAI-T that predicts fear learning in SCR responding (ß_st_ = −0.167), and the unique variance in IUS that predicts fear learning in startle responding (ß_st_ = −0.149), rather than the shared variance across the questionnaires/scales, and across outcome measures. Note that these standardized coefficients of the final model are identical to the coefficients estimated with the final path model in Step 2.

### Study 1: Interim discussion

As expected, we observed moderate to strong links between the scores derived from the three questionnaires/scales linked to negative emotionality (STAI-*T*, NEO-FFI-*N*, IUS) in a large sample, whereas links between the three outcome measures of conditioned responding (SCRs, startle, ratings) were weak to moderate. These weak to moderate correlations among outcome measures are consistent with the idea that they tap into slightly different processes and capture different timings with respect to CS processing (discussed in Lonsdorf & Merz, 2017). Uncorrected univariate analyses, which would mirror the approach employed in studies focusing on these questionnaires/scales and outcome measures in isolation, suggest significant associations between STAI-T and SCR, IUS and SCR, and IUS and startle responding. Additionally these univariate analyses suggest trends for the NEO-FFI-N and SCR and startle, as well as for STAI-T and startle responding. Importantly, the correlation coefficients between one outcome measure and the different questionnaire scores do not differ significantly from each other

The results from the multivariate path model on the other hand may suggest a slightly different conclusion. When all measures are integrated in one statistical model, only associations between STAI-T and CS discrimination in SCR, and between IUS and CS discrimination in startle responding remain significant. This may suggest a certain level of specificity indicating that more general measures (such as the STAI-T) of negative emotionality might be specifically associated with outcome measures that also reflect very general physiological arousal or general affective processes (SCR). Our results suggest that these may be distinct from more specific measures (such as the IUS) in the negative emotionality domain, which seem to be associated with measures reflecting valence specific processes related to fear learning.

Remarkably, the structural equation model that includes latent variables for negative emotionality and fear responding only, reveals a strong association between both latent variables with good fit. All questionnaires/scales load strongly on the latent factor negative emotionality, with STAI-T showing the strongest factor loading. These high factor loadings indicate that including negative emotionality as latent factor may be indeed appropriate and informative. The fear learning latent variable is represented by less strong, but still medium sized factor loadings, suggesting that these readouts are related and there might be a broad underlying “fear learning” variable, but it also underlines that there is room for dissociation between them in particular because from a theoretical and neurobiological perspective (Hamm & Weike, 2005) these different outcome measures capture slightly different cognitive-affective processes and timings with respect to CS processing (Leuchs et al., 2019; Lonsdorf et al., 2017; Lonsdorf, Klingelhöfer-Jens, Andreatta, Beckers, Chalkia, Gerlicher, Jentsch, et al., 2019). The initial effect between the latent variables is eliminated when adding the two identified specific paths.

In sum, our results speak in favor of a substantial shared variance and the existence of a latent “negative emotionality” factor across the three questionnaires and scales included, speaking in favor of convergent validity. In addition, specific paths between specific scales and outcome measures were identified - yet it cannot be excluded that these specific results may represent overfitting in this particular sample. In sum, our results may indicate that it might rather be the shared variance across the questionnaires/scales that predicts fear learning and the specificity of the here identified paths needs to be replicated and further investigated in another well-powered sample potentially including item-level factor analytical approaches which would require a substantially larger sample than included in this study.

### Study 2: Brief Introduction

Surprisingly, the neurocognitive processes underlying an association between negative emotionality and the ability to discriminate signals of danger and safety remain largely unknown to date (Lonsdorf & Merz, 2017). A bunch of studies has investigated neural associations with trait anxiety (Barrett & Armony, 2009; Indovina et al., 2011; Morriss et al., 2015; Sehlmeyer et al., 2011b; Tzschoppe et al., 2014), intolerance of uncertainty (Barrett & Armony, 2009; Indovina et al., 2011; Morriss et al., 2015; Sehlmeyer et al., 2011b; Tzschoppe et al., 2014) or neuroticism (Barrett & Armony, 2009; Indovina et al., 2011; Morriss et al., 2015; Sehlmeyer et al., 2011b; Tzschoppe et al., 2014), but studies integrating fMRI results with concurrently acquired psychophysiological measures and measures of negative emotionality in one and the same study are rare or even non-existent in the field (reviewed in Lonsdorf & Merz, 2017).

Therefore, we aim to address this fundamental gap in the literature and extend our findings from Study 1 by exploring the neuro-functional mechanisms potentially underlying the observed specific association between the STAI-*T* score with CS+/CS− discrimination in SCRs that has been observed in Study 1. To achieve this aim, we re-analyzed data from a large pre-existing sample (N = 113).

Of note, recording of startle responses in the MR scanner has been challenging due to technical challenges, which we have only very recently been able to overcome (Kuhn et al., 2019). To date, we do not have sufficiently large samples with simultaneous recordings of startle and BOLD-fMRI to follow up upon the association between the IUS and CS+/CS− discrimination in startle.

### Study 2: Methods

#### Participants

Study 2 is based on a pre-existing dataset. One-hundred and twenty four participants were included in Study 2. Participants were recruited from a large screening sample (described in Kuhn et al., 2015) in which they had been screened by a psychologist for neurological disorders using the M.I.N.I. interview (Sheehan et al., 1998) as a diagnostic tool. Any current or prior psychiatric neurological disorder or self-reported abuse of illegal drugs led to exclusion. Participants were re-invited for two separate studies based on exposure to life adversity in order to study its impact on a post-extinction manipulation (i.e., reinstatement; an experimental phase not included in the analyses of the current manuscript and published elsewhere (R Scharfenort et al., 2016), first study also in (Scharfenort and Lonsdorf, 2016), second unpublished). These two studies employed identical experimental protocols (including experimenter) for fear acquisition, and thus participants were pooled across both studies into Study 2 of this manuscript.

Out of the 124 participants, eleven participants had to be excluded due to technical issues (N = 6), pathological anatomy (N = 2) or missing items on the STAI-*T* (N = 3) which resulted in 113 participants included for fMRI, rating and SCR analyses of fear acquisition. The final sample included 44 females and 69 males, ranging in age between 19 and 34 years. On average participants were 25 ± 3.5 (SD) years old. All participants had normal or corrected to normal vision.

The studies were conducted in accordance with the Declaration of Helsinki and approved by the Ethical Review Board of the General Medical Council Hamburg. All participants gave written informed consent to participate. Participants received 50 Euros for their participation.

#### Questionnaires

Participants filled in a batch of questionnaires prior to the experiment. This batch included 1) questions to obtain demographic information, 2) the state scale of the State-Trait Anxiety Inventory (Spielberger et al., 1983), and 2) the NEO-FFI (Gerhard, 1999; McCrae & Costa Jr., 2004). The trait scale of the State-Trait Anxiety Inventory (STAI-*T*) was already acquired within the context of the screening sample, and based on the results obtained in Study 1, the STAI-*T* is of main interest here.

Overall reliability of the STAI-*T* was high in the final sample as indicated by Cronbach’s α = 0.93. Notably, a smaller range of STAI-T scores was obtained in Study 2 as compared to Study 1. STAI-T scores ranged between 20 and 59 with a mean of 35 ± 9 (SD).

After completing the experiment (i.e., after fear acquisition training in the MR-environment), participants filled in a post-experimental awareness questionnaire (Lonsdorf et al., 2017) and results were orally confirmed with the experimenter. Participants were asked to estimate the total number of received electrotactile and other experimental stimuli, as well as about perceived CS-US contingencies curing the experiment (first as free recall, then forced choice). Consequently, participants were classified as either aware (N = 101, able to correctly report CS-US contingencies in free recall and/or forced choice) or unaware (N = 12, unable to report correct CS-US contingencies across questions).

#### Instructions

As in study 1, participants were not instructed with respect to the CS-US contingencies or the learning element of the study.

#### Visual material

Two different white fractals presented on a grey background (RGB [230,230,230], 340×320 pixel, resolution: 1024×768) served as conditioned stimuli (CSs; duration: 6-8s, mean: 7s). A white cross in the middle of the grey background screen served as inter trial interval (ITI; duration: 10-16s; mean: 13s). One of the fractals (CS+) co-terminated with the unconditioned stimulus (US) during all fear acquisition training trials (100% reinforcement rate), whereas the other fractal did not (CS-). A 100% reinforcement rate was chosen to render it likely that all participants learn the association between CS+ and US within the 14 presentations. To allow for a differentiation between CS+ and US-related neural activity despite of this high reinforcement rate, the CS+ duration was jittered between 6-8sec (mean=7sec). Allocation of the fractals to CS+/CS− was counterbalanced over all subjects.

#### Electrotactile US

Similar to Study 1, the US consisted of a train of three electrotactile stimuli (interval 50ms, duration 10ms). The US was administered through a surface electrode on the dorsal part of the right hand via a DS7A electrical stimulator (Digitimer, Elwyn Garden City, UK). Before the experiment started, intensity was calibrated individually to a maximum tolerable level with a mean US intensity ± SD of 7.18 ± 4.47 mA, see study 1 for details on the US calibration procedure.

#### CS-US awareness

CS-US awareness was assessed as in Study 1 with the exception that the interview was conducted immediately after fear acquisition training and not at the end of the whole experiment. Participants were classified as aware (N=101) or unaware (N=12) of CS-US contingencies.

#### Procedure

Fear acquisition training occurred between 1-6 pm. As mentioned, experimental phases on the subsequent day (extinction, reinstatement and reinstatement test) are not of interest to the current manuscript. Participants were instructed outside the MR-environment. They were instructed to attend the visual stimuli on the screen, no instruction with respect to CS-US contingency was given.

After positioning the participant within the MR-environment, two skin conductance recording electrodes were attached, as well as one stimulation electrode for US delivery. When positioned in the MR-scanner, the US calibration procedure was started. Next the actual experiment started with a CS habituation phase. Both CSs were explicitly unreinforced and presented seven times each. In the subsequent uninstructed fear acquisition training phase of interest, both CSs were presented 14 times each.

Presentation of all stimuli was controlled using Presentation Software (NeuroBehavioral Systems, Albany California, USA). After completing the experiment for that day, thus after fear acquisition training, participants completed the post experimental awareness questionnaire.

#### Subjective data recording – Subjective Ratings

Subjective ratings were acquired retrospectively, i.e., after all fear acquisition training trials. Participants indicated their level of stress/fear/tension elicited by the preceding CS+ and CS-presentations on a 25-stepped visual analog scale (VAS, anchored at 0 and 100). Ratings of participants failing to confirm their rating for both CS types, were set to missing values (N = 9). In case participants missing either the CS+ or the CS-rating, the respective rating was replaced with the mean CS+ or CS-rating of all valid responses of the other participants on that rating (number of replaced values CS+ = 7, CS-= 4).

#### Physiological data recording and processing – Skin conductance responses

SCRs were recorded using a BIOPAC MP100 amplifier (BIOPAC Systems Inc., Goleta, California, USA). Ag/AgCl electrodes were placed on the palmar side of the left hand on the distal and proximal hypothenar. Data were processed and scored according to published guidelines (Boucsein *et al*, 2012). Skin conductance data were down-sampled to 10Hz. The phasic SCRs to the CS onsets were manually scored off-line, using custom-made software. SCR amplitudes (in µS) were scored as the first response initiating 0.9-4.0s after CS onset (Boucsein et al., 2012). To normalize the distribution, the SCRs were log transformed (Venables & Christie, 1980) and range-corrected by division through an individuals’ maximum response amplitude (Lykken & Venables, 1971).

#### BOLD-fMRI

MRI data were acquired on a 3-Tesla MR-scanner (MAGNETOM trio, Siemens, Germany) using a 32-channel head coil. Functional data were obtained using an echo planar images (EPI) sequence (TR=2460ms, TE=26ms). For each volume, 40 slices with a voxel size of 2×2×2 mm (1mm gap) were acquired sequentially. Structural images were obtained by using a T1 MPRAGE sequence. Preprocessing and analyses were performed using standard pre-procsessing in SPM8 (Welcome Trust Centre for Neuroimaging, UCL, London, UK). Preprocessing included, coregisteration to the individual structural image, realignment, normalization to group-specific templates (created via the DARTEL-algorithm; (Ashburner, 2007) as well as smoothing (6mm FWHM).

#### Statistical analyses

For fMRI data, four effects-of-interest regressors were built at the first level (i.e. early and late half of the acquisition trials for CS+ and CS-) as well as ten nuisance regressors (BOLD responses for CS+ and CS-of two habituation regressors, USs, ratings, and six movement parameters derived from realignment). All regressors of interest were modeled as stick function and time locked to stimulus onset for acquisition analyses. The general linear model was used to compute regression coefficients (beta values) for the regressors in each voxel.

CS discrimination contrasts (CS+>CS-; CS->CS+ for the full acquistion phase) were estimated on the first level and taken into the second level analysis employing voxel-wise regression analyses with the STAI-T. The main effect of task was estimated in a full factorial model with two regressors for both CS+ and CS-(early and late) and contrasts of interest were set to CS+>CS-and CS->CS+ covering the full phase.

ROI analyses were based on key areas for fear conditioning [primary ROI: amygdala; secondary ROIs: hippocampus, dorsal anterior cingulate cortex/dmPFC (dmPFC), pallidum/putamen, ventromedial prefrontal cortex (vmPFC), thalamus, insula (Fullana *et al*, 2015; Sehlmeyer *et al*, 2011a)]. Masks were derived from the Harvard-Oxford cortical and subcortical structural atlases; with a probability threshold of 0.7. As no vmPFC and dmPFC mask is provided by the Harvard-Oxford atlases, we used a box (20 × 16×16mm) centered on coordinates from a previous (independent) study [vmPFC: x,y,z: 0,40,-12; dmPFC: 0, 43, 29 (Lonsdorf et al., 2014)]. Due to strong *a-priori* predictions with respect to the amygdala and the use of additional regions as secondary ROIs, correction was performed separately for each ROI. A statistical threshold of p<0.05 (FWE corrected within the ROI) was considered significant.

Parameter estimates from peak voxels were extracted from the first individual level. Correlation test were performed for CS discrimination as well as CS+ and CS-specific responding in SCRs and ratings with scores on the STAI-T. For ratings and SCR correlational analyses were also carried out with the parameter estimates.

Correlation analyses and data visualization was performed in R version 3.6.0 (2019-04-26) using the dplyr, tydr, corrplot, cowplot, ggplot2, and pych packages.

### Study 2: Results

#### Main effects of task

Successful fear acquisition was evident by significantly larger SCR amplitudes for the CS+ than for the CS-during the full fear acquisition phase [t(112) = 10.14, p < 0.001, *d* = 0.95]. Similarly, post-acquisition fear ratings were higher for the CS+ as compared to the CS-[t(102) = 15.65, p < 0.001, *d* = 1.52]. On a neuro-functional level (CS-discrimination (CS+>CS-; Table 1) was reflected in areas typically activated in fear acquisition (i.e., thalamus, amygdala, dmPFC/dACC, insula/frontal operculum and putamen/pallidum). Stronger activation to the CS-than the CS+ was observed in the vmPFC (T-maps are available on neurovault https://identifiers.org/neurovault.image:305007).

**Table 1.** Neural activation reflecting CS+/CS− discrimination during fear acquisition (main effects of task) in the defined ROIs for left (L) and right (R) regions separately. Cluster size (k), MNI coordinates (x,y,z), and statistical values for uncorrected (0.001uc) and family-wise error corrected p-values in the ROI using small-volume correction (SVC_FWE_) are reported. Note that we used seven ROIs with the amygdala being the primary ROI and the other six being secondary ROIs. Correcting for multiple comparisons related to these seven regions would yield a corrected significance threshold of 0.007 (0.05/7), which the right and left hippocampus would not meet. Coordinates are in MNI space.

| Contrast | Brain area | k | x | y | z | T | Z | p(uc) | p(svc<sub>FWE</sub>) |
| --- | --- | --- | --- | --- | --- | --- | --- | --- | --- |
| CS+>CS- | dmACC | 691 | -4 | 28 | 26 | 10.57 | Inf | <0.001 | <0.001 |
|  | amygdala (R) | 148 | 20 | -4 | -12 | 6.32 | 6.18 | <0.001 | <0.001 |
|  | amygdala (L) | 85 | -20 | -4 | -12 | 6.22 | 5.09 | <0.001 | <0.001 |
|  | thalamus (R) | 585 | 8 | -18 | -2 | 11.11 | Inf | <0.001 | <0.001 |
|  | thalamus (L) | 685 | -8 | -18 | -2 | 11.94 | Inf | <0.001 | <0.001 |
|  | insula (R) | 327 | 34 | 22 | 2 | 14.27 | Inf | <0.001 | <0.001 |
|  | insula (L) | 332 | -32 | 20 | 4 | 14.41 | Inf | <0.001 | <0.001 |
|  | hippocampus (L) | 3 | -18 | -40 | 0 | 3.67 | 3.64 | <0.001 | 0.013 |
|  | hippocampus (L) | 4 | 34 | -16 | -14 | 3.58 | 3.55 | <0.001 | 0.018 |
|  |  | 4 | 28 | -8 | -28 | 3.58 | 3.55 | <0.001 | 0.017 |
|  |  | 11 | 26 | -28 | -8 | 3.41 | 3.39 | <0.001 | 0.029 |
|  | pallidum (L) | 225 | -12 | 2 | 0 | 10.29 | Inf | <0.001 | <0.001 |
|  | putamen (L) | 638 | -22 | 12 | -6 | 9.35 | Inf | <0.001 | <0.001 |
|  | pallidum (R) | 190 | 12 | 4 | -2 | 11.50 | Inf | <0.001 | <0.001 |
|  | putamen (R) | 556 | 22 | 12 | -8 | 8.45 | Inf | <0.001 | <0.001 |
|  | vmPFC | 52 | 16 | 24 | -20 | 5.96 | 5.84 | <0.001 | <0.001 |
| CS->CS+ | vmPFC | 712 | 0 | 24 | -24 | 5.28 | 5.20 | <0.001 | <0.001 |
|  |  | 11 | -14 | 56 | 4 | 4.36 | 4.31 | <0.001 | <0.001 |
|  | dmPFC | 156 | 20 | 28 | 46 | 5.42 | 5.34 | <0.001 | <0.001 |
|  |  | 430 | -12 | 38 | 42 | 5.15 | 5.08 | <0.001 | <0.001 |

#### Associations of CS+/CS− discrimination in SCRs and ratings with STAI-*T* scores

We explored associations between the STAI-*T* score and CS+/CS− discrimination in SCRs, post-experimental ratings as well as in BOLD-fMRI. In contrast to what was observed in Study 1, the STAI-*T* score was not significantly associated with CS+/CS− discrimination in SCRs or ratings during fear acquisition training in univariate correlation analyses (SCR: r = −0.05, p = 0.59; Rat: r = −0.15, p = 0.13) as illustrated in

Figure 4. Despite the absence of differences in CS discrimination we explored possible associations with CS+ or CS-responding individually. STAI-*T* was weakly (r = 0.31, p_BH_ < 0.01) and positively correlated with CS-responding (i.e., higher CS-ratings in individuals with higher STAI-T scores) in ratings, but not with either CS+ or CS-responding in SCR.

**Figure 4.**
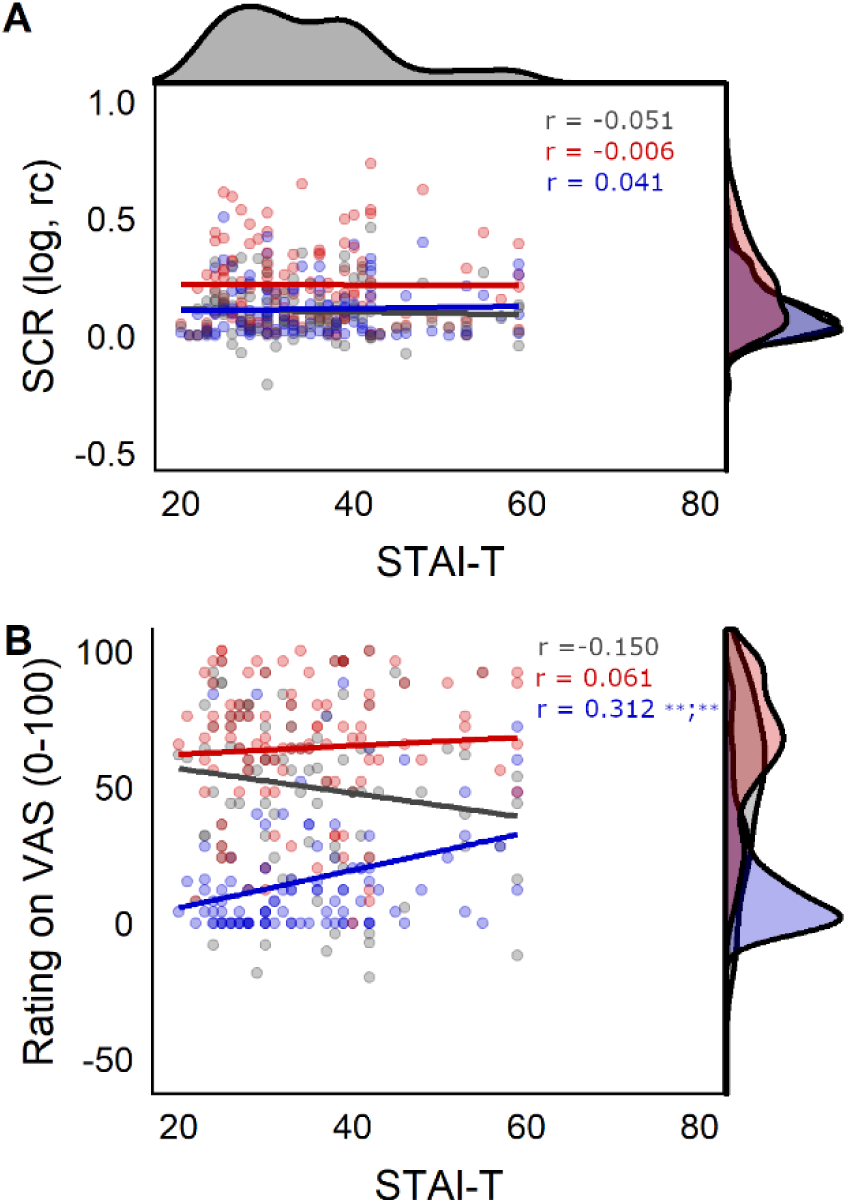
Scatterplots showing associations between (A) the STAI-T and SCR (skin conductance responses) and (B) between STAI-T and ratings for CS discrimination in grey, CS+ responding in red, and CS- responding in blue. Corresponding correlation coefficients (r) are displayed in corresponding colors. The density distribution of scores on the STAI-T is displayed on top of the figure, density distributions per stimulus type (CS+, CS-) and for CS discrimination are displayed on the right side of the figure for each dependent variable (SCR, ratings). ** indicates p < 0.01 for raw and Benjamini Hochberg (BH) adjusted p-values separated by a semicolon. Blank space indicates p > 0.1.

#### Neuro-functional associations of CS+/CS− discrimination with STAI-*T* scores

On a neural level, however, higher STAI-*T* scores were associated with significantly stronger CS+/CS− discrimination related activation of the right amygdala, the putamen (bilaterally) and the thalamus (bilaterally) during fear acquisition training (Table *2*, Figure 5). The corresponding T-map is available on Neurovault: https://identifiers.org/neurovault.image:305007). The association of the STAI-T scores seems to be driven by a positive association of the STAI-T scores with neural activation to the CS+ (right amygdala: x,y,z: 24,-10,-14; k=21, T:3.57, Z:3.46, pSVC(_FWE_): 0.008; left Putamen: x,y,z: −22,18,-2; k=11, T:3.59, Z:3.48, pSVC(_FWE_): 0.018; right Putamen: x,y,z: 22,20,-4; k=35, T:4.01, Z:3.87, pSVC(_FWE_): 0.005; left Thalamus: x,y,z: −4, −22, 14; k=178, T:4.71, Z:4.49, pSVC(_FWE_): 0.001; right Thalamus: x,y,z: 16, −30, 12; k=38, T:3.65, Z:3.4, pSVC(_FWE_): 0.001) but not the CS-(no significant effects in the ROIs).

**Figure 5.**
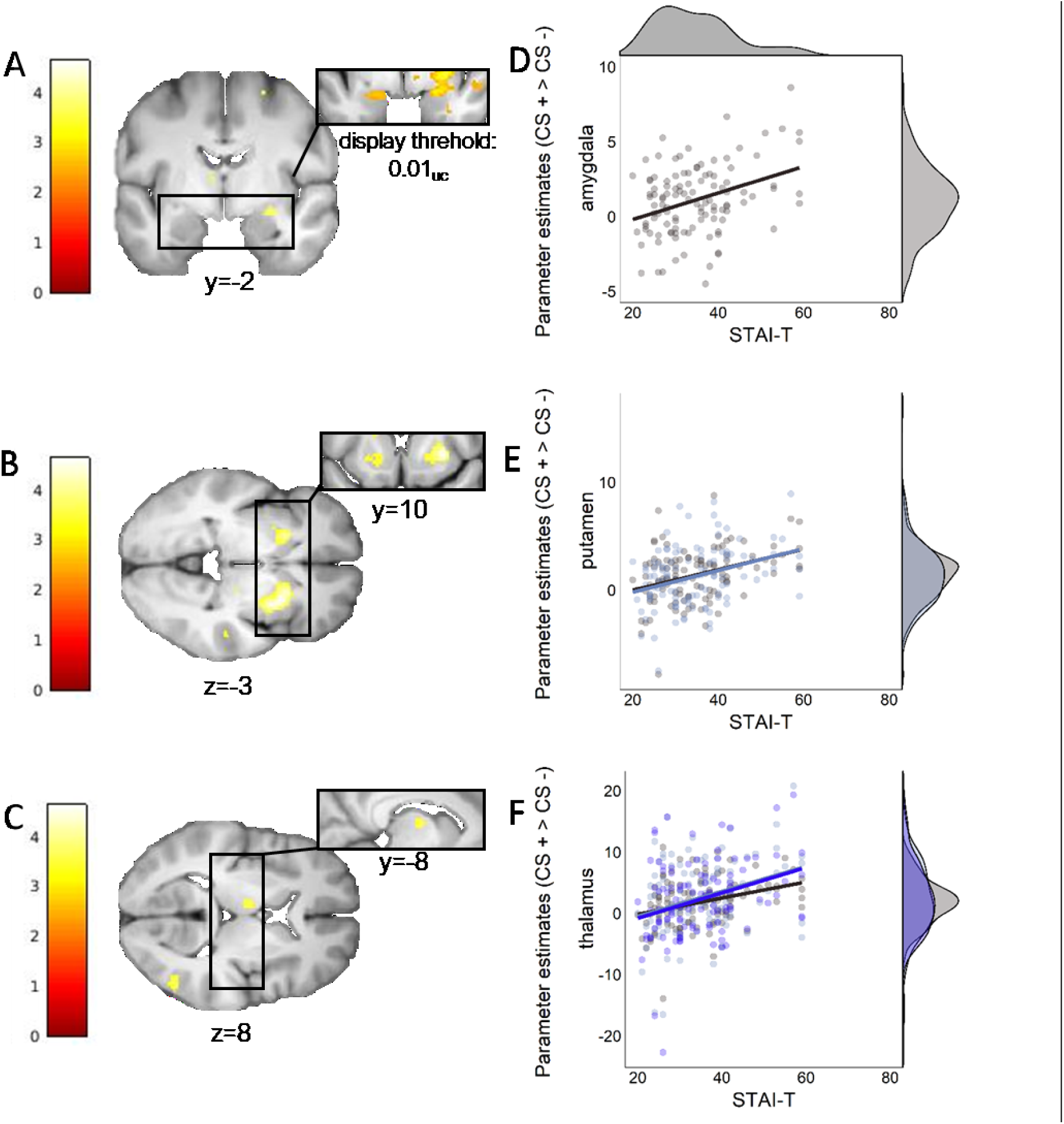
Neural activation for CS+/CS− discrimination (CS+>CS-) during fear acquisition training for areas significantly activated when regressed with STAI-T for the (A) amygdala [26, −10, −12], (B) putamen [28, 12, −2; −22, 16, −4], and (C) thalamus [(−8, −8, 8; −2); (−20, 12; 4), (4, −20, 12)] on the left. Coordinates are in MNI space. Scatterplots on the right represent the association between STAI-T scores and extracted peak voxel parameter estimates. Note that for the putamen and thalamus multiple peak values per ROI are displayed in different shades. Density distributions for peak value estimates are shown on the right, density distribution of STAI-T values in Study 2 is displayed on top of the upper scatterplot. A display threshold of uncorrected (uc) p < 0.001 was used to illustrate the extent of peak activations, unless otherwise specified. Note that statistics are based on SVC_FWE_ –corrected values.

**Table 2.** Neural activation reflecting significant ROI-based results (p<0.05 SVCFWE) for a regression of STAI-*T* on CS discrimination (CS+>CS-) during fear acquisition training. Cluster size k and coordinates x, y and z of the respective cluster are reported. Note that we used seven ROIs with the amygdala being the primary ROI and the other six being secondary ROIs. Correcting for multiple comparisons related to these seven regions would yield a corrected threshold of 0.007 (0.05/7), which the right and left thalamus would not meet for the full STAI-*T*. Coordinates are in MNI space.

| **STAI-*T*** |  |  |  |  |  |  |  |  |  |
| --- | --- | --- | --- | --- | --- | --- | --- | --- | --- |
| **Contrast** | **Brain area** | **k** | **x** | **y** | **z** | **T** | **Z** | **p(uc)** | **p(SVC<sub>FWE</sub>)** |
| **CS+>CS-** | putamen (R) | 230 | 28 | 12 | -2 | 4.56 | 4.36 | <0.001 | **0.001** |
|  | putamen (L) | 75 | -22 | 16 | -4 | 4.06 | 3.91 | <0.001 | **0.005** |
|  | amygdala (R) | 13 | 26 | -10 | -12 | 3.95 | 3.81 | <0.001 | **0.003** |
|  | thalamus (L) | 32 | -8 | -8 | 8 | 3.60 | 3.50 | <0.001 | **0.023** |
|  |  | 10 | -2 | -20 | 12 | 3.50 | 3.40 | <0.001 | **0.023** |
|  | thalamus (R) | 12 | 4 | -20 | 12 | 3.47 | 3.38 | <0.001 | **0.033** |
| **CS->CS+** | none |  |  |  |  |  |  |  |  |

Robustness checks revealed that a model including the covariate ‘life adversity’ (as participants in Study 2 were initially recruited based on this variable) yielded comparable results. More precisely, statistical values differed only at the last decimal place with the exception of the right thalamus which does not meet the 0.05 threshold when including the covariate ‘life adversity’ (data not shown). Of note, these areas are also significantly implicated in CS-discrimination irrespective of STAI-*T* in this sample (see above).

Of these areas, CS discrimination in SCRs was only positively associated with peak voxel activation in the left putamen (r = 0.240, p_BH_ = 0.028), and with the first cluster in the left thalamus (r = 0.322, p_BH_ = 0.002). The latter might be driven by a positive association between SCRs for the CS+ and the left thalamus (r = 0.315, p_BH_ = 0.002), whereas the CS-is does not show a significant association. A graphical representation of these associations can be found in the Supplementary Material (**Supplemental Figure 1**). Ratings for CS discrimination, the CS+ and the CS-were not significantly associated with peak voxel activation in any of our ROIs (all p_BH_ > 0.512).

#### Exploratory analyses testing for an association with the STAI-*T* score and US intensity as well as awareness

Finally, as previous research has suggested an association between awareness and US intensity with trait-anxiety (for a review see Lonsdorf & Merz, 2017) exploratory analyses were conducted for study 2 to exclude that results are biased by these factors. In brief, neither awareness of CS contingencies nor US intensity were significantly associated with the STAI-*T*: US intensity in mA did not correlate with STAI-*T* scores (r = −0.11, *p* = 0.24). Individuals aware (n = 101, mean STAI-*T* = 35) and unaware (n = 12, mean STAI-*T* = 32) of CS-US contingencies did not differ significantly in STAI-*T* scores (Welch two sample t-test: t(20.5) = 1.64, *p* = 0.16).

#### Comparing the STAI-T distributions across both samples (behavioral study vs. fMRI study)

To explore whether the distribution of STAI-T values in Study 1 and Study 2 are different (see Figure 6A), a two-sample Kolmogorov-Smirnov test was performed. This test indicates that both samples come from different distributions, D =. 219, p <. 001. As can be derived from Figure 6B the fMRI sample includes substantially more individuals with low STAI-T values (i.e., < 50) as compared to the behavioral sample.

**Figure 6.**
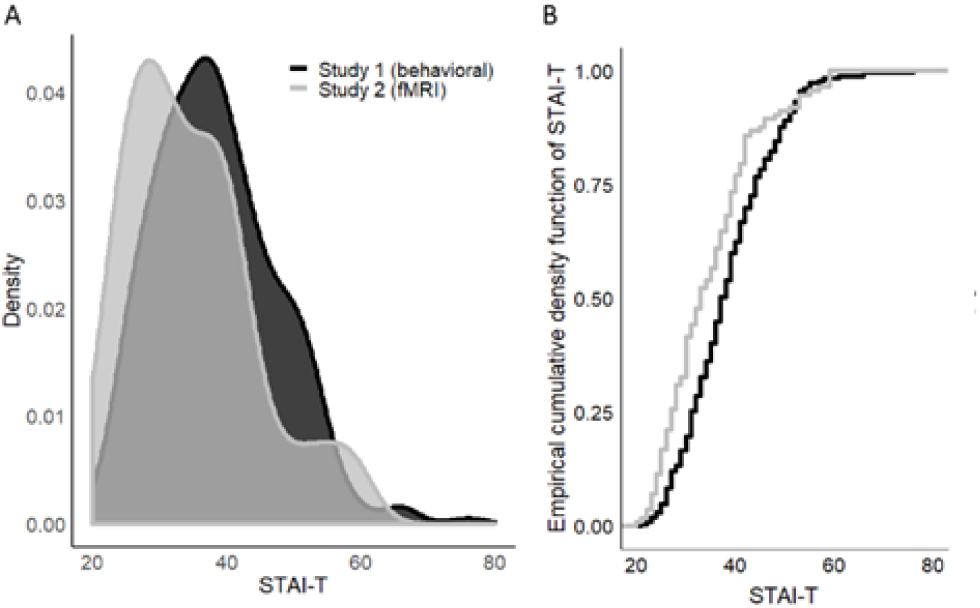
Distribution of STAI-T scores in the sample included in behavioral Study 1 (black) and fMRI Study 2 (grey) illustrated as (A) overlapping densities for both samples and (B) the empirical cumulative density function for both samples.

### Study 2: Interim summary

Results of study 2 show a positive association of the STAI-*T* score with CS+/CS− discrimination on a neuro-functional level in the right amygdala, the putamen (bilateral) and the thalamus (bilateral). These regions have all been implicated in the ability to discriminate signals of danger from signals of safety, both in the literature (Fullana et al., 2015a; Sehlmeyer C et al., 2009) and in the paradigm and sample reported here. Of note, the amygdala is a core region implicated in fear learning (Büchel et al., 1998; Greco & Liberzon, 2016; Herry & Johansen, 2014; LaBar et al., 1998; Tovote et al., 2015) and has been previously linked to individual differences in discriminating signals of danger from signals of safety (Indovina et al., 2011). This previous work has often not included the simultaneous acquisition of both autonomic (i.e., SCRs) and neuro-functional measures in the same experimental phase (Furmark et al., 1997; MacNamara et al., 2015) while others have recoded both measures during fear acquisition (Barrett & Armony, 2009; Sehlmeyer et al., 2011b) or fear expression (Indovina et al., 2011). Importantly, also in domains of threat processing, similar positive associations between STAI-*T* score and amygdala reactivity as reported here have been observed (Ewbank et al., 2009; Fonzo et al., 2015; Michely et al., 2020; Stout et al., 2017). In addition, our work provides evidence for an involvement of the amygdala in individual differences underlying the *strength of fear learning* beyond the average (i.e., a general role in fear acquisition and expression). This is important as evidence suggesting the role of the amygdala in fear acquisition has been questioned (Fullana et al., 2015b) and also as there is accumulating evidence that aggregated results across a group do not necessarily generalize to individuals (e.g., Fisher et al., 2018; Hedge et al., 2017).

Despite the observed associations with the STAI-*T* score and CS+/CS− discrimination on a neural level, we did not observe a significant association between CS+/CS− discrimination in SCRs and the STAI-*T* score as observed in Study 1. Yet, the sample in Study 2 (N=113) is substantially smaller than the sample in Study 1 (N=356). Calculating the required sample size to determine if the correlation coefficient of -.183 – as observed between STAI-T and CS+/CS− discrimination in Study 1 – does significantly differ from zero yields a required sample of 232 individuals (assuming an α of 0.05 and a power (1-β) of 80%; http://www.sample-size.net/correlation-sample-size/). This is twice the number of subjects included in Study 2 and hence the non-replication of SCR results should be treated with caution. Furthermore, we highlight that the distribution of STAI-*T* scores between Study 1 and Study 2 (see Figure 6) is significantly different. The distribution in Study 2 is substantially more left skewed. More precisely study 1 (behavioral) contained more individuals with a high STAI-*T* score (> 60) than Study 2 (fMRI) – as evident from the much flatter right tail of the density in Study 2. In Study 1 scores reach values up to 76, whereas in Study 2 the maximum score is 59. Moreover, in the imaging study (Study 2) there are proportionally more individuals included with STAI-T scores falling in the lower quartile and thus in a group that would be characterized as having no or low anxiety (STAI-T < 37). Hence, we call for caution when interpreting this null finding as a replication failure of findings in Study 1. Instead, sample bias – possibly originating from high anxious individuals not signing up for fMRI studies – may in addition to the differences in power between the studies also contribute to different results in both studies. Hence, we replicate a recent report of the existence of a profound sampling bias in MRI studies in a large set of pooled studies, which showed that participants in MRI studies had lower trait anxiety scores compared to participants in behavioral studies (Charpentier et al., 2020). This implies that good characterization and reporting of study populations and experimental parameters is highly important especially in individual difference research (Lonsdorf & Merz, 2017).

#### General discussion Study 1 and 2

The overarching aim of this work was to investigate and explore the contribution of putatively specific and shared variance between three commonly used questionnaires related to negative emotionality on conditioned responding across multiple units of analysis (ratings, skin conductance, startle, BOLD-fMRI) in a fear conditioning paradigm in two large samples (N_Study1_=356; N_Study2_=113).

The three questionnaires selected for this study (the trait scale of the STAI, the neuroticism scale of the NEO-FFI and the Intolerance of Uncertainty Scale) were selected because of the abundance of literature in the field of individual differences in fear conditioning research (for a review Lonsdorf & Merz, 2017). These three questionnaires share a substantial part of their variance, here operationalized as a latent ‘negative emotionality’ variable. Our results hint to potentially specific associations between the STAI-T (Study 1), with the discrimination between cues signaling danger (CS+) or safety (CS-) in the arousal-related outcome measure of skin conductance responding, and intolerance of uncertainty with discriminating danger and safety in valence related outcome measures of startle responses. These results should be interpreted, however, with caution as overfitting in this particularly study cannot be excluded. Importantly, we also find support for the existence of a negative emotionality latent variable that has an effect on general fear learning.

Notably, *not* accounting for shared variance between measures of emotional negativity in univariate correlational analyses revealed comparable negative associations of *all three* questionnaires with CS+/CS− discrimination in SCRs - with the STAI-T showing the strongest and significant association. Note that association with NEO-FFI N and IUS were not statistically significant but correlation coefficients were comparable to the one derived from the STAI-T/SCR association.

Results derived from the multivariate path model approach imply that the observed univariate associations of NEO-FFI-N and IUS with SCR CS+/CS− discrimination might be fully explained by their shared variance with STAI-T. Of note, the observed association between the STAI-T and CS+/CS− discrimination was negative in Study 1 and Study 2 (i.e., high scores are associated with less discrimination) – although clearly non-significant in Study 2 - while others have observed positive associations in small samples (Indovina et al., 2011; MacNamara et al., 2015). There is a plethora of potential and plausible reasons underlying these seeming discrepancies. As discussed in a recent review (Lonsdorf & Merz, 2017), these include a number of procedural factors including high vs. low reinforcement rate (Dunsmoor et al., 2007), potency of the experimental situation (Shmuel Lissek et al., 2006) instructions (Hugdahl & Ohman, 1977), additional triggered outcome measures such as startle that impact on the learning process (Sjouwerman et al., 2016b) as well as sample biases or exclusion of specific participants (Charpentier et al., 2020; Lonsdorf, Klingelhöfer-Jens, Andreatta, Beckers, Chalkia, Gerlicher, Haaker, et al., 2019) to name just a few. Hence, it is possible that neither results may be necessarily ‘wrong’ as different associations may unfold depending on the specific sample, experimental context or boundary conditions. While the large sample in Study 1 – in comparison to the typical sample size in the field in individual difference studies (systematically summarized in Lonsdorf & Merz, 2017) - should contribute to trust in our findings, systematic investigations are highly warranted. Hence, we urge authors to focus more on procedural details, demands and related processes, and potential sampling bias in future studies to explore whether this may facilitate mechanistic conclusions (Lonsdorf & Merz, 2017).

Of note, in the substantially smaller Study 2, we do, not observe an association between the STAI-T score with CS+/CS− discrimination in SCRs or ratings. Our sample size calculation revealed that Study 2 was most likely underpowered to detect an association between CS+/CS− discrimination an STAI-T scores (given the correlation coefficient observed in Study 1) and in addition represents individuals sampled from a different distribution, likely caused by the nature of the study (i.e., fMRI). Hence, we replicate the recent results by a report suggesting samples for fMRI and behavioral studies are drawn from different populations (Charpentier et al., 2020).

Given that nearly all published studies in the literature (with few exceptions (J. Haaker et al., 2015)) fall well below the sample size in Study 1 (N=356), the zero findings across outcome-measures in these studies (Arnaudova et al., 2013; Barrett & Armony, 2009; Chin, Nelson, Jackson, et al., 2016; Joos et al., 2012; Sehlmeyer et al., 2011b; Torrents-Rodas et al., 2013) are difficult to interpret. More large-N studies are needed to determine whether these different results originate from fluctuations around the null (i.e., absence of a true effect) or whether there is indeed a true effect.

In Study 2, we did observe a positive association between the STAI-T and CS-responding in subjective ratings (i.e., higher CS-ratings in individuals with higher scores). This may be in line with the non-significant positive trend observed in univariate analyses for CS-ratings and STAI-*T* in Study 1. Note that for all three questionnaires/scales in Study 1 (i.e, STAI-T, NEO-FFI-N, IUS) the association between the respective questionnaire/scale and CS-responding in ratings was trend wise significant. A link between negative emotionality and CS-responding is in line with the suggestion of deficient safety signal processing in individuals with affective disorders or those at risk (Craske et al., 2014); previous results in a similarly large sample (J. Haaker et al., 2015); as well as previous reports on associations between STAI-T and deficits in *safety signal* (e.g., CS-) processing (Gazendam et al., 2013; J. Haaker et al., 2015; Haddad et al., 2012; Kindt & Soeter, 2014). Results should however be treated with caution, as this association was only observed in the smaller Study 2, and was only a trend in Study 1.

In Study 2, however, CS+/CS− discrimination in a number of brain areas of key relevance to fear processing and expression are positively associated with the STAI-T score as well as its sub-components (amygdala, bilateral putamen, bilateral thalamus). The construct of trait anxiety as assessed by the STAI-T has been criticized in the literature for representing a psychometrically inhomogeneous scale (Reiss, 1997), capturing facets of both anxiety and depression (Bados et al., 2010; Bieling et al., 1998; Mathews et al., 1996; Reiss, 1997). Factor analyses on the single items of these questionnaires in larger well powered studies could address this question further. Exploratory factor analyses performed on the items included in Study 1 are not included here as the results likely represented over-fitting in this particular study sample (i.e., items derived from one scale loaded primarily on a single factor and few plausibly expected cross loadings between similar items across scales were observed). Note that the sample size of Study 1 in relation to the number of items was too small for this purpose and hence, data are not shown and included. Future work in appropriately sized samples should focus on unraveling cross-questionnaire factors that may inform us on the specific mechanisms and components underlying the association with negative emotionality and the discrimination between danger and safety cues (CS+/CS− discrimination) in fear acquisition.

It is also noteworthy that the selection of questionnaires related to negative emotionality for study 1 was exclusively motivated by evidence from the available literature in the field of human fear conditioning (Lonsdorf & Merz, 2017), in which the three selected measures have commonly been used (in isolation however). Hence future work should extend these findings by targeting additional measures not included in this report, such as specific measures of depression, to further unravel an underlying potentially mechanistic component of negative emotionality driving the link between dispositional negativity and fear learning on an autonomic level.

Furthermore, it is noteworthy, that we observe a specific association not only between differential SCRs and the STAI-T score but also between fear potentiated startle (i.e., CS+>CS-in startle responding) and intolerance of uncertainty scores. Although awaiting replication of this potentially interesting finding, it its noteworthy that others have also observed intolerance of uncertainty scores to be *negatively* associated with startle responding during the uncertain but not certain threat condition (Nelson & Shankman, 2011) suggesting that it was not predictive of general aversive responding, but specific to responses to uncertain averseness.

Importantly, despite our work providing clear evidence for substantially shared variance between the three questionnaires, the specific dissociations in outcome measures and questionnaire scores (i.e., specific association of STAI-T with CS-discrimination in SCRs, and IUS with CS-discrimination in FPS) may provide insights into the underlying processes. Different outcome measures capture and reflect diverse aspects and represent unique sources of variance in fear processing (Lonsdorf et al., 2017) and emotional processing per se (Lang et al., 1993; Mauss & Robinson, 2009). SCRs are thought to reflect general arousal. Startle in turn is considered a rather fear specific index (Lonsdorf et al., 2017) that per definition reflects an enhanced reflexive response towards an unexpected, and therewith uncertain, event. Hence, both results may carry complementary mechanistic information corresponding to multi-causal vulnerability in fear and anxiety. As it was technically not yet feasible to implement combined EMG-fMRI measurements at the time of data acquisition, future studies profiting from this novel option (Kuhn et al., 2019; Lindner et al., 2015) are warranted to investigate the neurobiological mechanisms underlying the specific association between intolerance of uncertainty and FPS.

Our results clearly highlight the value of multimodal work and multivariate analyses tools and suggest that ‘compound profiles’ that integrate multiple input and outcome measures and hence potentially capture multiple processes may in the long run prove useful from a ‘personalized medicine’ perspective. Yet, the associations observed here between measures of negative emotionality and physiological responding are not substantially large or even of medium size. This has to be kept in mind when discussing implications for their potential for biomarker development. Yet, we speculate that a multivariate composite of different response pattern (‘profile’) may show stronger associations which may hold potential for the development of clinically useful products in the long run.

In sum, it is fundamental to uncover factors and their potential interactions that contribute to individual risk and resilience to pathological fear – although fear conditioning protocols may rather model adaptive fear (Beckers et al., 2013). Hence, improved understanding of the personality related and neurobiological processes underlying individual differences in experimental fear learning can be expected to translate into improved understanding on how adaptive responding to threats turns into maladaptive fear responding (Jovanovic & Ressler, 2010; Ressler & Mayberg, 2007). It will thus be important to extend the investigation of individual differences and the underlying neurobiology beyond experimental fear acquisition to tests focusing on the long-term retention of fear and extinction memory (i.e., return of fear (Martínez et al., 2012)), and ultimately to clinical populations. We provide a very first step towards this overarching aim towards ultimateleyimproving our mechanistic understanding of pathological fear and emotional responding by providing initial insights of inter-individual differences in fear processing using multivariate approaches across units of analysis in two samples.

The authors declare no conflict of interest.

## References

Allan, T. A., & DeYoung, C. G. (2016). Personality Neuroscience and the Five Factor Model. In The Oxford Handbook of the Five Factor Model. Oxford University Press.

Allen, T. A., & DeYoung, C. G. (2017). Personality Neuroscience and the Five Factor Model. The Oxford Handbook of the Five Factor Model. https://doi.org/10.1093/oxfordhb/9780199352487.013.26

Altman, D. G., & Royston, P. (2006). The cost of dichotomising continuous variables. BMJ: British Medical Journal, 332(7549), 1080.

Arnaudova, I., Krypotos, A.-M., Effting, M., Boddez, Y., Kindt, M., & Beckers, T. (2013). Individual differences in discriminatory fear learning under conditions of ambiguity: A vulnerability factor for anxiety disorders? Personality and Social Psychology, 4, 298. https://doi.org/10.3389/fpsyg.2013.00298

Ashburner, J. (2007). A fast diffeomorphic image registration algorithm. NeuroImage, 38(1), 95–113. https://doi.org/10.1016/j.neuroimage.2007.07.007

Bados, A., Gómez-Benito, J., & Balaguer, G. (2010). The state-trait anxiety inventory, trait version: Does it really measure anxiety? Journal of Personality Assessment, 92(6), 560–567. https://doi.org/10.1080/00223891.2010.513295

Balsamo, M., Romanelli, R., Innamorati, M., Ciccarese, G., Carlucci, L., & Saggino, A. (2013). The State-Trait Anxiety Inventory: Shadows and Lights on its Construct Validity. Journal of Psychopathology and Behavioral Assessment, 35(4), 475–486. https://doi.org/10.1007/s10862-013-9354-5

Barlow, D. H., Sauer-Zavala, S., Carl, J. R., Bullis, J. R., & Ellard, K. K. (2014). The Nature, Diagnosis, and Treatment of Neuroticism: Back to the Future. Clinical Psychological Science, 2(3), 344–365. https://doi.org/10.1177/2167702613505532

Barrett, J., & Armony, J. L. (2009). Influence of trait anxiety on brain activity during the acquisition and extinction of aversive conditioning. Psychological Medicine, 39(2), 255–265. https://doi.org/10.1017/S0033291708003516

Beckers, T., Krypotos, A.-M., Boddez, Y., Effting, M., & Kindt, M. (2013). What’s wrong with fear conditioning? Biological Psychology, 92(1), 90–96. https://doi.org/10.1016/j.biopsycho.2011.12.015

Bieling, P. J., Antony, M. M., & Swinson, R. P. (1998). The State-Trait Anxiety Inventory, Trait version: Structure and content re-examined. Behaviour Research and Therapy, 36(7-8), 777–788.

Blumenthal, T. D., Cuthbert, B. N., Filion, D. L., Hackley, S., Lipp, O. V., & Van Boxtel, A. (2005). Committee report: Guidelines for human startle eyeblink electromyographic studies. Psychophysiology, 42, 1–15. https://doi.org/10.1111/j.1469-8986.2005.00271.x

Boswell, J. F., Thompson-Hollands, J., Farchione, T. J., & Barlow, D. H. (2013). Intolerance of Uncertainty: A Common Factor in the Treatment of Emotional Disorders. Journal of Clinical Psychology, 69(6). https://doi.org/10.1002/jclp.21965

Boucsein, W., Fowles, D. C., Grimnes, S., Ben-Shakhar, G., Roth, W. T., Dawson, M. E., & Filion, D. L. (2012). Publication recommendations for electrodermal measurements. Psychophysiology, 49, 1017–1034. https://doi.org/10.1111/j.1469-8986.2012.01384.x

Büchel, C., Morris, J., Dolan, R. J., & Friston, K. J. (1998). Brain systems mediating aversive conditioning: An event-related fMRI study. Neuron, 20(5), 947–957.

Bufferd, S. J., Dougherty, L. R., Olino, T. M., Dyson, M. W., Carlson, G. A., & Klein, D. N. (2018). Temperament Distinguishes Persistent/Recurrent from Remitting Anxiety Disorders Across Early Childhood. Journal of Clinical Child and Adolescent Psychology: The Official Journal for the Society of Clinical Child and Adolescent Psychology, American Psychological Association, Division 53, 47(6), 1004–1013. https://doi.org/10.1080/15374416.2016.1212362

Buhr, K., & Dugas, M. J. (2002). The Intolerance of Uncertainty Scale: Psychometric properties of the English version. Behav Res Ther, 40(8), 931–945. https://doi.org/10.1016/S0005-7967(01)00092-4

Charpentier, C. J., Faulkner, P., pool, eva, Ly, V., Tollenaar, M. S., Kluen, L. M., Fransen, A., Yamamori, Y., Lally, N., Mkrtchian, A., Valton, V., Huys, Q. J., Morrow, K., Krenz, V., Kalbe, F., Cremer, A., Zerbes, G., Kausche, F. M., Wanke, N., … O’Doherty, J. (2020). How representative are neuroimaging samples? Large-scale evidence for trait anxiety differences between MRI and behaviour-only research participants. [Preprint]. PsyArXiv. https://doi.org/10.31234/osf.io/cqdne

Chin, B., Nelson, B. D., Chin, B., Nelson, B. D., Jackson, F., & Hajcak, G. (2016). Intolerance of Uncertainty and Startle Potentiation in in Relation to Different Threat Reinforcement Rates. International Journal of Psychophysiology, 99(April 2016), 79–84. https://doi.org/10.1016/j.ijpsycho.2015.11.006

Chin, B., Nelson, B. D., Jackson, F., & Hajcak, G. (2016). Intolerance of uncertainty and startle potentiation in relation to different threat reinforcement rates. International Journal of Psychophysiology: Official Journal of the International Organization of Psychophysiology, 99, 79–84. https://doi.org/10.1016/j.ijpsycho.2015.11.006

Cohen, J. (1983). The Cost of Dichotomization. Applied Psychological Measurement, 7(3), 249–253. https://doi.org/10.1177/014662168300700301

Craske, M. G., Treanor, M., Conway, C. C., Zbozinek, T., & Vervliet, B. (2014). Maximizing exposure therapy: An inhibitory learning approach. Behaviour Research and Therapy, 58, 10–23. https://doi.org/10.1016/j.brat.2014.04.006

Dawson, M. E., Schell, A. M., Filion, D. L., & Berntson, G. G. (2007). The Electrodermal System. In Handbook of Psychophysiology (Third edition). Cambridge University Press. http://dx.doi.org/10.1017/CBO9780511546396.007

DeYoung, C. G. (2015). Cybernetic Big Five Theory. Journal of Research in Personality, 56, 33–58. https://doi.org/10.1016/j.jrp.2014.07.004

Dugas, M. J., Gagnon, F., Ladouceur, R., & Freeston, M. H. (1998). Generalized anxiety disorder: A preliminary test of a conceptual model. Behaviour Research and Therapy, 36(2), 215–226.

Duits, P., Cath, D. C., Lissek, S., Hox, J. J., Hamm, A. O., Engelhard, I. M., van den Hout, M. A., & Baas, J. M. P. (2015). Updated meta-analysis of classical fear conditioning in the anxiety disorders. 32, 239–253. https://doi.org/10.1002/da.22353

Dunsmoor, J. E., Bandettini, P. A., & Knight, D. C. (2007). Impact of continuous versus intermittent CS-UCS pairing on human brain activation during Pavlovian fear conditioning. Behavioral Neuroscience, 121(4), 635–642. https://doi.org/10.1037/0735-7044.121.4.635

Dunsmoor, J. E., Campese, V. D., Ceceli, A. O., Ledoux, J. E., & Phelps, E. A. (2016). Novelty-facilitated extinction: Providing a novel outcome in place of an expected threat diminishes recovery of defensive responses. Biological Psychiatry, 78(3), 203–209. https://doi.org/10.1016/j.biopsych.2014.12.008.Novelty-facilitated

Ewbank, M. P., Lawrence, A. D., Passamonti, L., Keane, J., Peers, P. V., & Calder, A. J. (2009). Anxiety predicts a differential neural response to attended and unattended facial signals of anger and fear. NeuroImage, 44(3), 1144–1151. https://doi.org/10.1016/j.neuroimage.2008.09.056

Eysenck, H. J. (1950). Dimensions of Personality. Transaction Publishers.

Fergus, T. A. (2013). A comparison of three self-report measures of intolerance of uncertainty: An examination of structure and incremental explanatory power in a community sample. Psychological Assessment, 25(4), 1322–1331. https://doi.org/10.1037/a0034103

Fisher, A. J., Medaglia, J. D., & Jeronimus, B. F. (2018). Lack of group-to-individual generalizability is a threat to human subjects research. Proceedings of the National Academy of Sciences, 115(27), E6106–E6115. https://doi.org/10.1073/pnas.1711978115

Fonzo, G. A., Ramsawh, H. J., Flagan, T. M., Sullivan, S. G., Letamendi, A., Simmons, A. N., Paulus, M. P., & Stein, M. B. (2015). Common and disorder-specific neural responses to emotional faces in generalised anxiety, social anxiety and panic disorders. The British Journal of Psychiatry: The Journal of Mental Science, 206(3), 206–215. https://doi.org/10.1192/bjp.bp.114.149880

Forcadell, E., Torrents-Rodas, D., Vervliet, B., Leiva, D., Tortella-Feliu, M., & Fullana, M. A. (2017). Does fear extinction in the laboratory predict outcomes of exposure therapy? A treatment analog study. International Journal of Psychophysiology: Official Journal of the International Organization of Psychophysiology, 121, 63–71. https://doi.org/10.1016/j.ijpsycho.2017.09.001

Freeston, M. H., Rhéaume, J., Letarte, H., Dugas, M. J., & Ladouceur, R. (1994). Why do people worry? Personality and Individual Differences, 17(6), 791–802. https://doi.org/10.1016/0191-8869(94)90048-5

Fullana, M. A., Harrison, B. J., Soriano-Mas, C., Vervliet, B., Cardoner, N., Àvila-Parcet, A., & Radua, J. (2015a). Neural signatures of human fear conditioning: An updated and extended meta-analysis of fMRI studies. Molecular Psychiatry. https://doi.org/10.1038/mp.2015.88

Fullana, M. A., Harrison, B. J., Soriano-Mas, C., Vervliet, B., Cardoner, N., Àvila-Parcet, A., & Radua, J. (2015b). Neural signatures of human fear conditioning: An updated and extended meta-analysis of fMRI studies. Molecular Psychiatry. https://doi.org/10.1038/mp.2015.88

Furmark, T., Fischer, H., Wik, G., Larsson, M., & Fredrikson, M. (1997). The amygdala and individual differences in human fear conditioning. Neuroreport, 8(18), 3957–3960.

Gazendam, F. J., Kamphuis, J. H., & Kindt, M. (2013). Deficient safety learning characterizes high trait anxious individuals. Biological Psychology, 92(2), 342–352. https://doi.org/10.1016/j.biopsycho.2012.11.006

Gerhard, U. (1999). Borkenau, P. & Ostendorf, F. (1993). NEO-Fünf-Faktoren Inventar (NEO-FFI) nach Costa und McCrae. Göttingen: Hogrefe. Preis DM 84.-. Zeitschrift Für Klinische Psychologie Und Psychotherapie, 28(2), 145–146. https://doi.org/10.1026//0084-5345.28.2.145

Gerlach, A. L., Andor, T., & Patzelt, J. (2008a). Die Bedeutung von Unsicherheitsintoleranz für die Generalisierte Angststörung Modellüberlegungen und Entwicklung einer deutschen Version der Unsicherheitsintoleranz-Skala. Zeitschrift Für Klinische Psychologie Und Psychotherapie, 37(3), 190–199. https://doi.org/10.1026/1616-3443.37.3.190

Gerlach, A. L., Andor, T., & Patzelt, J. (2008b). Die Bedeutung von Unsicherheitsintoleranz für die Generalisierte Angststörung Modellüberlegungen und Entwicklung einer deutschen Version der Unsicherheitsintoleranz-Skala. Zeitschrift Für Klinische Psychologie Und Psychotherapie, 37(3), 190–199. https://doi.org/10.1026/1616-3443.37.3.190

Gerlach, A. L., Andor, T., & Patzelt, J. (2008c). Die Bedeutung von Unsicherheitsintoleranz für die Generalisierte Angststörung Modellüberlegungen und Entwicklung einer deutschen Version der Unsicherheitsintoleranz-Skala. Zeitschrift Für Klinische Psychologie Und Psychotherapie, 37(3), 190–199. https://doi.org/10.1026/1616-3443.37.3.190

Goldstein, B. L., Kotov, R., Perlman, G., Watson, D., & Klein, D. N. (2018). Trait and facet-level predictors of first-onset depressive and anxiety disorders in a community sample of adolescent girls. Psychological Medicine, 48(8), 1282–1290. https://doi.org/10.1017/S0033291717002719

Greco, J. A., & Liberzon, I. (2016). Neuroimaging of Fear-Associated Learning. Neuropsychopharmacology: Official Publication of the American College of Neuropsychopharmacology, 41(1), 320–334. https://doi.org/10.1038/npp.2015.255

Grupe, D. W., & Nitschke, J. B. (2013). Uncertainty and anticipation in anxiety: An integrated neurobiological and psychological perspective. Nature Reviews Neuroscience, 14(7), 488–501. https://doi.org/10.1038/nrn3524

Guo, W., Xue, J.-M., Shao, D., Long, Z.-T., & Cao, F.-L. (2015). Effect of the interplay between trauma severity and trait neuroticism on posttraumatic stress disorder symptoms among adolescents exposed to a pipeline explosion. PloS One, 10(3), e0120493. https://doi.org/10.1371/journal.pone.0120493

Haaker, J., Lonsdorf, T. B., Schümann, D., Menz, M., Brassen, S., Bunzeck, N., Gamer, M., & Kalisch, R. (2015). Deficient inhibitory processing in trait anxiety: Evidence from context-dependent fear learning, extinction recall and renewal. Biological Psychology, 111, 65–72. https://doi.org/10.1016/j.biopsycho.2015.07.010

Haaker, Jan, Maren, S., Andreatta, M., Merz, C. J., Richter, J., Richter, S. H., Drexler, S. M., Lange, M. D., Jüngling, K., Nees, F., Seidenbecher, T., Fullana, M. A., Wotjak, C. T., & Lonsdorf, T. B. (2019). Making translation work: Harmonizing cross-species methodology in the behavioural neuroscience of Pavlovian fear conditioning. Neuroscience & Biobehavioral Reviews. https://doi.org/10.1016/j.neubiorev.2019.09.020

Haddad, A. D. M., Pritchett, D., Lissek, S., & Lau, J. Y. F. (2012). Trait anxiety and fear responses to safety cues: Stimulus generalization or sensitization? Journal of Psychopathology and Behavioral Assessment, 34, 323–331. https://doi.org/10.1007/s10862-012-9284-7

Hakulinen, C., Elovainio, M., Pulkki-Råback, L., Virtanen, M., Kivimäki, M., & Jokela, M. (2015). Personality and Depressive Symptoms: Individual-Participant Meta-Analysis of 10 Cohort Studies. Depression and Anxiety, 32(7), 461–470. https://doi.org/10.1002/da.22376

Hamm, A. O., & Weike, A. I. (2005). The neuropsychology of fear learning and fear regulation. International Journal of Psychophysiology: Official Journal of the International Organization of Psychophysiology, 57(1), 5–14. https://doi.org/10.1016/j.ijpsycho.2005.01.006

Hedge, C., Powell, G., & Sumner, P. (2017). The reliability paradox: Why robust cognitive tasks do not produce reliable individual differences. Behavior Research Methods, 1–21. https://doi.org/10.3758/s13428-017-0935-1

Hengartner, M. P., Ajdacic-Gross, V., Wyss, C., Angst, J., & Rössler, W. (2016). Relationship between personality and psychopathology in a longitudinal community study: A test of the predisposition model. Psychological Medicine, 46(8), 1693–1705. https://doi.org/10.1017/S0033291716000210

Hengartner, Michael P., Tyrer, P., Ajdacic-Gross, V., Angst, J., & Rössler, W. (2018). Articulation and testing of a personality-centred model of psychopathology: Evidence from a longitudinal community study over 30 years. European Archives of Psychiatry and Clinical Neuroscience, 268(5), 443–454. https://doi.org/10.1007/s00406-017-0796-8

Hengartner, Michael P., van der Linden, D., Bohleber, L., & von Wyl, A. (2017). Big Five Personality Traits and the General Factor of Personality as Moderators of Stress and Coping Reactions Following an Emergency Alarm on a Swiss University Campus. Stress and Health: Journal of the International Society for the Investigation of Stress, 33(1), 35–44. https://doi.org/10.1002/smi.2671

Herry, C., & Johansen, J. P. (2014). Encoding of fear learning and memory in distributed neuronal circuits. Nature Neuroscience, 17(12), 1644–1654. https://doi.org/10.1038/nn.3869

Holmes, A. J., & Patrick, L. M. (2018). The Myth of Optimality in Clinical Neuroscience. Trends in Cognitive Sciences, 22(3), 241–257. https://doi.org/10.1016/j.tics.2017.12.006

Hugdahl, K., & Ohman, A. (1977). Effects of instruction on acquisition and extinction of electrodermal responses to fear-relevant stimuli. Journal of Experimental Psychology. Human Learning and Memory, 3(5), 608–618.

Indovina, I., Robbins, T. W., N????ez-Elizalde, A. O., Dunn, B. D., & Bishop, S. J. (2011). Fear-Conditioning Mechanisms Associated with Trait Vulnerability to Anxiety in Humans. Neuron, 69(3), 563–571. https://doi.org/10.1016/j.neuron.2010.12.034

Jeronimus, B. F., Kotov, R., Riese, H., & Ormel, J. (2016). Neuroticism’s prospective association with mental disorders halves after adjustment for baseline symptoms and psychiatric history, but the adjusted association hardly decays with time: A meta-analysis on 59 longitudinal/prospective studies with 443 313 participants. Psychological Medicine, 46(14), 2883–2906. https://doi.org/10.1017/S0033291716001653

Joos, E., Vansteenwegen, D., & Hermans, D. (2012). Worry as a predictor of fear acquisition in a nonclinical sample. Behavior Modification, 36(5), 723–750. https://doi.org/10.1177/0145445512446477

Jovanovic, T., & Ressler, K. J. (2010). How the neurocircuitry and genetics of fear inhibition may inform our understanding of PTSD. The American Journal of Psychiatry, 167(6), 648–662. https://doi.org/10.1176/appi.ajp.2009.09071074

Julian, L. J. (2011). Measures of anxiety: State-Trait Anxiety Inventory (STAI), Beck Anxiety Inventory (BAI), and Hospital Anxiety and Depression Scale-Anxiety (HADS-A). Arthritis Care & Research, 63 Suppl 11, S467–472. https://doi.org/10.1002/acr.20561

Kalokerinos, E. K., Murphy, S. C., Koval, P., Bailen, N. H., Crombez, G., Hollenstein, T., Gleeson, J., Thompson, R. J., Ryckeghem, D. M. L. V., Kuppens, P., & Bastian, B. (2020). Neuroticism may not reflect emotional variability. Proceedings of the National Academy of Sciences. https://doi.org/10.1073/pnas.1919934117

Kindt, M., & Soeter, M. (2014). Fear Inhibition in High Trait Anxiety. 9(1). https://doi.org/10.1371/journal.pone.0086462

Krampen, G. (1981). IPC-Fragebogen zu Kontrollüberzeugungen. Verlag für Psychologie.

Kuhn, M., Scharfenort, R., Schümann, D., Schiele, M. A., Münsterkötter, A. L., Deckert, J., Domschke, K., Haaker, J., Kalisch, R., Pauli, P., Reif, A., Romanos, M., Zwanzger, P., & Lonsdorf, T. B. (2015). Mismatch or allostatic load? Timing of life-adversity differentially shapes gray matter volume and anxious-temperament. Social Cognitive and Affective Neuroscience, nsv137. https://doi.org/10.1093/scan/nsv137

Kuhn, M., Wendt, J., Sjouwerman, R., Büchel, C., Hamm, A., & Lonsdorf, T. B. (2019). The neurofunctional basis of affective startle modulation in humans – evidence from combined facial electromyography and functional magnetic resonance imaging. Biological Psychiatry, 0(0). https://doi.org/10.1016/j.biopsych.2019.07.028

LaBar, K. S., Gatenby, J. C., Gore, J. C., LeDoux, J. E., & Phelps, E. A. (1998). Human amygdala activation during conditioned fear acquisition and extinction: A mixed-trial fMRI study. Neuron, 20(5), 937–945.

Lahey, B. B. (2009). Public Health Significance of Neuroticism. The American Psychologist, 64(4), 241–256. https://doi.org/10.1037/a0015309

Lang, P. J., Bradley, M. M., & Cuthbert, B. N. (1990). Emotion, attention, and the startle reflex. Psychological Review, 97(3), 377–395.

Lang, P. J., Greenwald, M. K., Bradley, M. M., & Hamm, A. O. (1993). Looking at pictures: Affective, facial, visceral, and behavioral reactions. Psychophysiology, 30(3), 261–273. https://doi.org/10.1111/j.1469-8986.1993.tb03352.x

Lee, I.A., P., K. J. (2013). Calculation for the test of the difference between two dependent correlations with one variable in common. http://quantpsy.org.

Leuchs, L., Schneider, M., & Spoormaker, V. I. (2019). Measuring the conditioned response: A comparison of pupillometry, skin conductance, and startle electromyography. Psychophysiology, 56(1), e13283. https://doi.org/10.1111/psyp.13283

Lindner, K., Neubert, J., Pfannmöller, J., Lotze, M., Hamm, A. O., & Wendt, J. (2015). Fear-potentiated startle processing in humans: Parallel fMRI and orbicularis EMG assessment during cue conditioning and extinction. International Journal of Psychophysiology, May, 1–11. https://doi.org/10.1016/j.ijpsycho.2015.02.025

Lissek, S, Powers, A., McClure, E., Phelps, E., Woldehawariat, G., Grillon, C., & Pine, D. (2005). Classical fear conditioning in the anxiety disorders: A meta-analysis. BEHAVIOUR RESEARCH AND THERAPY, 43(11), 1391–1424. https://doi.org/10.1016/j.brat.2004.10.007

Lissek, Shmuel, Pine, D. S., & Grillon, C. (2006). The strong situation: A potential impediment to studying the psychobiology and pharmacology of anxiety disorders. Biological Psychology, 72(3), 265–270. https://doi.org/10.1016/j.biopsycho.2005.11.004

Lonsdorf, T. B., Haaker, J., & Kalisch, R. (2014). Long-term expression of human contextual fear and extinction memories involves amygdala, hippocampus and ventromedial prefrontal cortex: A reinstatement study in two independent samples. Social Cognitive and Affective Neuroscience, nsu018-. https://doi.org/10.1093/scan/nsu018

Lonsdorf, T. B., Klingelhöfer-Jens, M., Andreatta, M., Beckers, T., Chalkia, A., Gerlicher, A., Haaker, J., Jentsch, V., Mertens, G., Drexler, S. M., Richter, J., Sjouwerman, R., Wendt, J., & Merz, C. J. (2019). How to not get lost in the garden of forking paths: Lessons learned from human fear conditioning research regarding exclusion criteria [Preprint]. PsyArXiv. https://doi.org/10.31234/osf.io/6m72g

Lonsdorf, T. B., Klingelhöfer-Jens, M., Andreatta, M., Beckers, T., Chalkia, A., Gerlicher, A., Jentsch, V. L., Meir Drexler, S., Mertens, G., Richter, J., Sjouwerman, R., Wendt, J., & Merz, C. J. (2019). Navigating the garden of forking paths for data exclusions in fear conditioning research. ELife, 8. https://doi.org/10.7554/eLife.52465

Lonsdorf, T. B., Menz, M. M., Andreatta, M., Fullana, M. A., Golkar, A., Haaker, J., Heitland, I., Hermann, A., Kuhn, M., Kruse, O., Drexler, S. M., Meulders, A., Nees, F., Pittig, A., Richter, J., Römer, S., Shiban, Y., Schmitz, A., Straube, B., … Merz, C. J. (2017). Don’t fear “fear conditioning”: Methodological considerations for the design and analysis of studies on human fear acquisition, extinction, and return of fear. Neuroscience and Biobehavioral Reviews. https://doi.org/10.1016/j.neubiorev.2017.02.026

Lonsdorf, T. B., & Merz, C. J. (2017). More than just noise: Inter-individual differences in fear acquisition, extinction and return of fear in humans—Biological, experiential, temperamental factors, and methodological pitfalls. Neuroscience and Biobehavioral Reviews, 80(April), 703–728. https://doi.org/10.1016/j.neubiorev.2017.07.007

Lykken, D., & Venables, P. (1971). Direct measurement of skin conductance: A proposal for standardization. Psychophysiology, 8(5), 656–672. https://doi.org/10.1111/j.1469-8986.1971.tb00501.x

M. W. Browne, R. C. (1992). Alternative ways of assessing model fit. Sociological Methods & Research, 21(2). https://doi.org/10.1177/0049124192021002005

MacCallum, R. C., Browne, M. W., & Sugawara, H. M. (1996). Power analysis and determination of sample size for covariance structure modeling. Psychological Methods, 1(2), 130–149. https://doi.org/10.1037/1082-989X.1.2.130

MacNamara, A., Rabinak, C. A., Fitzgerald, D. A., Zhou, X. J., Shankman, S. A., Milad, M. R., & Phan, K. L. (2015). Neural correlates of individual differences in fear learning. Behavioural Brain Research, 287, 34–41. https://doi.org/10.1016/j.bbr.2015.03.035

Martínez, K. G., Castro-Couch, M., Franco-Chaves, J. A., Ojeda-Arce, B., Segura, G., Milad, M. R., & Quirk, G. J. (2012). Correlations between psychological tests and physiological responses during fear conditioning and renewal. Biology of Mood & Anxiety Disorders, 2(1), 16. https://doi.org/10.1186/2045-5380-2-16

Mathews, A., Ridgeway, V., & Williamson, D. A. (1996). Evidence for attention to threatening stimuli in depression. Behaviour Research and Therapy, 34(9), 695–705.

Mauss, I. B., & Robinson, M. D. (2009). Measures of emotion: A review. Cognition & Emotion, 23(2), 209–237. https://doi.org/10.1080/02699930802204677

McClelland, G., & Irwin, J. R. (2003). Negative Consequences of Dichotomizing Continuous Predictor Variables (SSRN Scholarly Paper ID 627741). Social Science Research Network. https://papers.ssrn.com/abstract=627741

McCrae, R. R., & Costa Jr., P. T. (2004). A contemplated revision of the NEO Five-Factor Inventory. Personality and Individual Differences, 36(3), 587–596. https://doi.org/10.1016/S0191-8869(03)00118-1

McEvoy, P. M., & Mahoney, A. E. J. (2012). To Be Sure, To Be Sure: Intolerance of Uncertainty Mediates Symptoms of Various Anxiety Disorders and Depression. Behavior Therapy, 43(3), 533–545. https://doi.org/10.1016/j.beth.2011.02.007

Michely, J., Rigoli, F., Rutledge, R. B., Hauser, T. U., & Dolan, R. J. (2020). Distinct Processing of Aversive Experience in Amygdala Subregions. Biological Psychiatry: Cognitive Neuroscience and Neuroimaging, 5(3), 291–300. https://doi.org/10.1016/j.bpsc.2019.07.008

Mineka, S., & Oehlberg, K. (2008). The relevance of recent developments in classical conditioning to understanding the etiology and maintenance of anxiety disorders. Acta Psychologica, 127(3), 567–580. https://doi.org/10.1016/j.actpsy.2007.11.007

Morriss, J., Christakou, A., & van Reekum, C. M. (2015). Intolerance of uncertainty predicts fear extinction in amygdala-ventromedial prefrontal cortical circuitry. Biology of Mood & Anxiety Disorders, 5. https://doi.org/10.1186/s13587-015-0019-8

Morriss, J., Christakou, A., & van Reekum, C. M. (2016). Nothing is safe: Intolerance of uncertainty is associated with compromised fear extinction learning. Biological Psychology. https://doi.org/10.1016/j.biopsycho.2016.05.001

Nelson, B. D., & Shankman, S. A. (2011). Does Intolerance of Uncertainty Predict Anticipatory Startle Responses to Uncertain Threat? International Journal of Psychophysiology: Official Journal of the International Organization of Psychophysiology, 81(2), 107–115. https://doi.org/10.1016/j.ijpsycho.2011.05.003

Otto, M. W., Leyro, T. M., Christian, K., Deveney, C. M., Reese, H., Pollack, M. H., & Orr, S. P. (2007). Prediction of “fear” acquisition in healthy control participants in a de novo fear-conditioning paradigm. Behavior Modification, 31(1), 32–51. https://doi.org/10.1177/0145445506295054

Preacher, K. J., Rucker, D. D., MacCallum, R. C., & Nicewander, W. A. (2005). Use of the extreme groups approach: A critical reexamination and new recommendations. Psychological Methods, 10(2), 178–192. https://doi.org/10.1037/1082-989X.10.2.178

Reiss, S. (1997). Trait anxiety: It’s not what you think it is. Journal of Anxiety Disorders, 11(2), 201–214.

Ressler, K. J., & Mayberg, H. S. (2007). Targeting abnormal neural circuits in mood and anxiety disorders: From the laboratory to the clinic. Nature Neuroscience, 10(9), 1116–1124. https://doi.org/10.1038/nn1944

Saulnier, K. G., Allan, N. P., Raines, A. M., & Schmidt, N. B. (2019). Depression and Intolerance of Uncertainty: Relations between Uncertainty Subfactors and Depression Dimensions. Psychiatry, 82(1), 72–79. https://doi.org/10.1080/00332747.2018.1560583

Scharfenort, R, Menz, M., & Lonsdorf, T. B. (2016). Adversity-induced relapse of fear: Neural mechanisms and implications for relapse prevention from a study on experimentally induced return-of-fear following fear conditioning and extinction. 6(7), e858–8. https://doi.org/10.1038/tp.2016.126

Scharfenort, Robert, & Lonsdorf, T. B. (2016). Neural correlates of and processes underlying generalized and differential return of fear. Social Cognitive and Affective Neuroscience, 612–620. https://doi.org/10.1093/scan/nsv142

Scheveneels, S., Boddez, Y., Vervliet, B., & Hermans, D. (2016). The validity of laboratory-based treatment research: Bridging the gap between fear extinction and exposure treatment. Behaviour Research and Therapy, 86, 87–94. https://doi.org/10.1016/j.brat.2016.08.015

Sehlmeyer, C., Dannlowski, U., Schöning, S., Kugel, H., Pyka, M., Pfleiderer, B., Zwitserlood, P., Schiffbauer, H., Heindel, W., Arolt, V., & Konrad, C. (2011a). Neural correlates of trait anxiety in fear extinction. Psychological Medicine, 41(4), 789–798. https://doi.org/10.1017/S0033291710001248

Sehlmeyer, C., Dannlowski, U., Schöning, S., Kugel, H., Pyka, M., Pfleiderer, B., Zwitserlood, P., Schiffbauer, H., Heindel, W., Arolt, V., & Konrad, C. (2011b). Neural correlates of trait anxiety in fear extinction. Psychological Medicine, 41(4), 789–798. https://doi.org/10.1017/S0033291710001248

Sehlmeyer C, Schoning S, Zwitserlood P, Pfleiderer B, Kircher T, & Arolt V. (2009). Human fear conditioning and extinction in neuroimaging: A systematic review. PLoS One, 4, e5865. https://doi.org/10.1371/journal.pone.0005865

Shackman, A. J., & Fox, A. S. (2018). Getting Serious about Variation: Lessons for Clinical Neuroscience (A Commentary on ’The Myth of Optimality in Clinical Neuroscience’). Trends in Cognitive Sciences, 22(5), 368–369. https://doi.org/10.1016/j.tics.2018.02.009

Shackman, A. J., Stockbridge, M. D., Tillman, R. M., Kaplan, C. M., Tromp, D. P. M., Fox, A. S., & Gamer, M. (2016). The neurobiology of dispositional negativity and attentional biases to threat: Implications for understanding anxiety disorders in adults and youth. Journal of Experimental Psychopathology, 7(3), 311–342. https://doi.org/10.5127/jep.054015

Shackman, A. J., Tromp, D. P. M., Stockbridge, M. D., Kaplan, C. M., Tillman, R. M., & Fox, A. S. (2016). Dispositional negativity: An integrative psychological and neurobiological perspective. Psychological Bulletin, 142(12), 1275–1314. https://doi.org/10.1037/bul0000073

Shackman, A. J., Weinstein, J. S., Hudja, S. N., Bloomer, C. D., Barstead, M. G., Fox, A. S., & Lemay, E. P. (2018). Dispositional negativity in the wild: Social environment governs momentary emotional experience. Emotion (Washington, D.C.), 18(5), 707–724. https://doi.org/10.1037/emo0000339

Sheehan, D. V., Lecrubier, Y., Sheehan, K. H., Amorim, P., Janavs, J., Weiller, E., Hergueta, T., Baker, R., & Dunbar, G. C. (1998). The Mini-International Neuropsychiatric Interview (M.I.N.I.): The development and validation of a structured diagnostic psychiatric interview for DSM-IV and ICD-10. In Journal of Clinical Psychiatry (Vol. 59, Issue SUPPL. 20, pp. 22–33). https://doi.org/10.1016/S0924-9338(99)80239-9

Sjouwerman, R., & Lonsdorf, T. B. (2018). Latency of skin conductance responses across stimulus modalities. Psychophysiology, October, 1–11. https://doi.org/10.1111/psyp.13307

Sjouwerman, R., Niehaus, J., Kuhn, M., & Lonsdorf, T. B. (2016a). Don ’ t startle me—Interference of startle probe presentations and intermittent ratings with fear acquisition. Psychophysiology, 53, 1889–1899. https://doi.org/10.1111/psyp.12761

Sjouwerman, R., Niehaus, J., Kuhn, M., & Lonsdorf, T. B. (2016b). Don’t startle me—Interference of startle probe presentations and intermittent ratings with fear acquisition. Psychophysiology, n/a-n/a. https://doi.org/10.1111/psyp.12761

Sjouwerman, R., Niehaus, J., & Lonsdorf, T. B. (2015). Contextual Change After Fear Acquisition Affects Conditioned Responding and the Time Course of Extinction Learning—Implications for Renewal Research. Frontiers in Behavioral Neuroscience, 9(December), 1–9. https://doi.org/10.3389/fnbeh.2015.00337

Spielberger, C. D., Gorsuch, R. L., & Lushene, R. E. (1983). Manual for the State-Trait Anxiety Inventory. Consulting Psychologists Press.

Spinhoven, P., Batelaan, N., Rhebergen, D., van Balkom, A., Schoevers, R., & Penninx, B. W. (2016). Prediction of 6-yr symptom course trajectories of anxiety disorders by diagnostic, clinical and psychological variables. Journal of Anxiety Disorders, 44, 92–101. https://doi.org/10.1016/j.janxdis.2016.10.011

Steunenberg, B., Beekman, A. T. F., Deeg, D. J. H., & Kerkhof, A. J. F. M. (2010). Personality predicts recurrence of late-life depression. Journal of Affective Disorders, 123(1-3), 164–172. https://doi.org/10.1016/j.jad.2009.08.002

Stout, D. M., Shackman, A. J., Pedersen, W. S., Miskovich, T. A., & Larson, C. L. (2017). Neural circuitry governing anxious individuals’ mis-allocation of working memory to threat. Scientific Reports, 7(1). https://doi.org/10.1038/s41598-017-08443-7

Torrents-Rodas, D., Fullana, M. A., Bonillo, A., Caseras, X., Andión, O., & Torrubia, R. (2013). No effect of trait anxiety on differential fear conditioning or fear generalization. Biological Psychology, 92(2), 185–190. https://doi.org/10.1016/j.biopsycho.2012.10.006

Tovote, P., Fadok, J. P., & Lüthi, A. (2015). Neuronal circuits for fear and anxiety. Nature Reviews Neuroscience, 16(6), 317–331. https://doi.org/10.1038/nrn3945

Tzschoppe, J., Nees, F., Banaschewski, T., Barker, G. J., Büchel, C., Conrod, P. J., Garavan, H., Heinz, A., Loth, E., Mann, K., Martinot, J.-L., Smolka, M. N., Gallinat, J., Ströhle, A., Struve, M., Rietschel, M., Schumann, G., Flor, H., & IMAGEN consortium. (2014). Aversive learning in adolescents: Modulation by amygdala-prefrontal and amygdala-hippocampal connectivity and neuroticism. Neuropsychopharmacology: Official Publication of the American College of Neuropsychopharmacology, 39(4), 875–884. https://doi.org/10.1038/npp.2013.287

Venables, P., & Christie, M. (1980). Electrodermal activity. In I. Martin & P. Venables (Eds.), Techniques in Pychophysiology (pp. 3–67). Chichester: Wiley.

Wichstrøm, L., Penelo, E., Rensvik Viddal, K., de la Osa, N., & Ezpeleta, L. (2018). Explaining the relationship between temperament and symptoms of psychiatric disorders from preschool to middle childhood: Hybrid fixed and random effects models of Norwegian and Spanish children. Journal of Child Psychology and Psychiatry, and Allied Disciplines, 59(3), 285–295. https://doi.org/10.1111/jcpp.12772

