## Supplemental Figure 1 for "Individual differences in fear acquisition: Multivariate analyses of different Emotional Negativity scales, physiological responding, subjective measures, and neural activation"

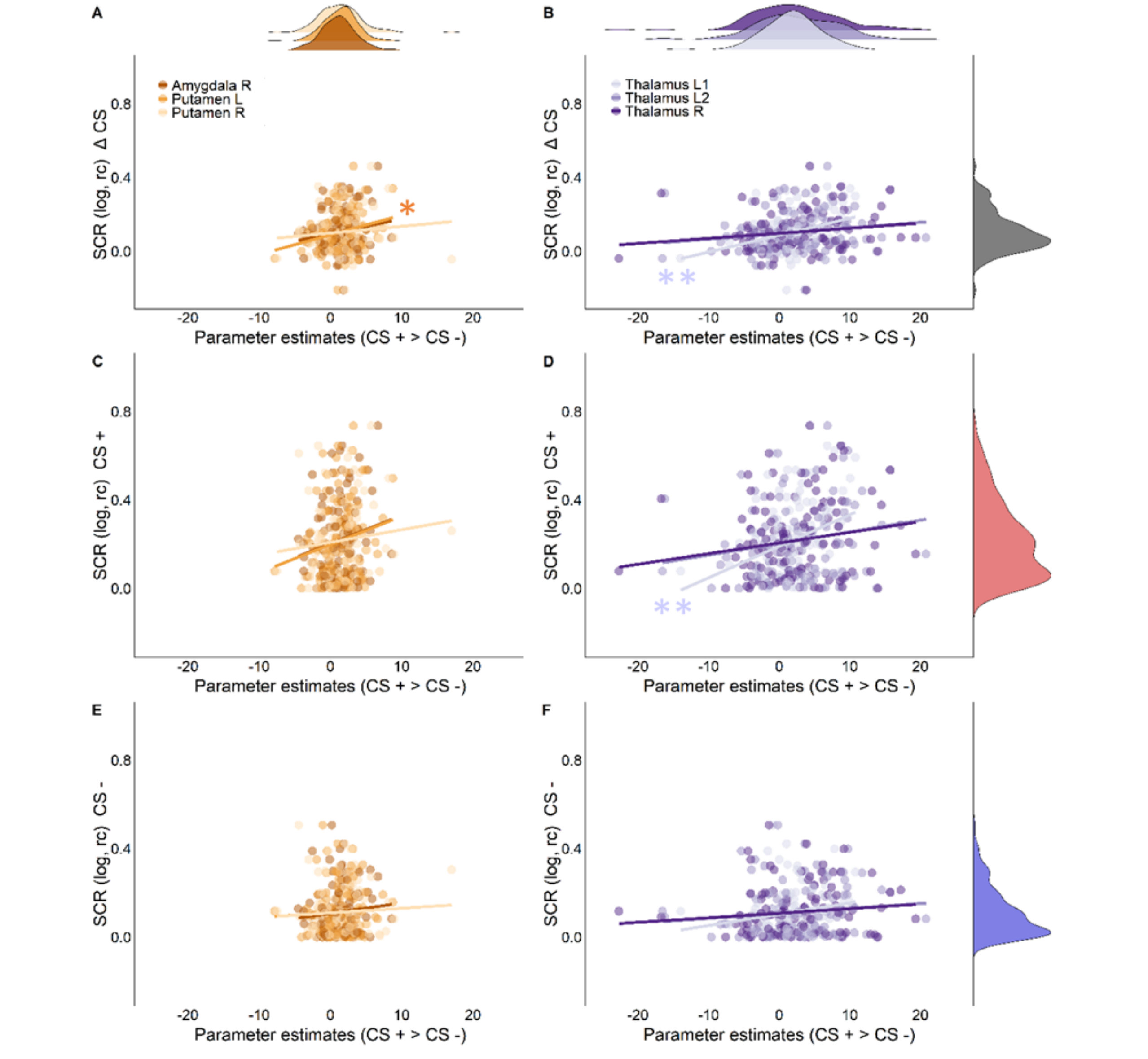


**Supplementary Figure 1**. Scatterplots for (A,B) CS+/CS- discrimination (C,D) CS+, and (E,F) CS- in SCR responding and parameter estimates for CS + > CS – contrasts in the six brain regions significantly associated with the STAI-T (parameter estimates extracted from peak voxels). The left panel shows the (A,C,E) right amygdala, left and right putamen and the right panel shows (B, D, E) two clusters in the left thalamus, and the right thalamus. Density distributions are shown on top of the figure for the parameter estimates (regions are color coded). Densities on the right side of each plot show SCR for CS discrimination, CS+ responding, and CS- responding respectively. * indicates $p_{BH} < 0.05$, ** indicates $p_{BH} < 0.01$.
